# Timely Schwann cell division during migration drives peripheral myelination *in vivo* via Laminin/cAMP pathway

**DOI:** 10.1101/2022.02.11.480035

**Authors:** Aya Mikdache, Marie-José Boueid, Emilie Lesport, Brigitte Delespierre, Julien Loisel-Duwattez, Cindy Degerny, Marcel Tawk

**Affiliations:** U1195, Inserm, University Paris-Saclay, 94276 Le Kremlin Bicêtre, France

**Keywords:** *sil*, Schwann cells, peripheral nervous system, mitotic spindle, MCPH, zebrafish, myelin, PLLn, mitosis, Laminin, cAMP

## Abstract

Schwann cells (SC) migrate along peripheral axons and divide intensively to generate the right number of cells prior to axonal ensheathment; however, little is known regarding the temporal and molecular control of their division, particularly during migration, and its impact on myelination. We report that Sil, a spindle pole protein associated with autosomal recessive primary microcephaly (MCPH), is required for temporal mitotic exit of SC. In *sil*-deficient *cassiopeia* (*csp^-/-^*) mutants, SC fail to radially sort and myelinate peripheral axons. Elevation of cAMP, but not Rac1 activity in *csp*^-/-^ restores myelin ensheathment. Most importantly, we show a significant decrease in Laminin expression within *csp^-/-^* posterior lateral line nerve and that forcing Laminin2 expression in *csp^-/-^* fully restores SC ability to myelinate. We also discovered that SC have a restricted time window during which they have to divide, while migrating, in order to trigger myelination. Thus, we unravel a novel and essential role for timely SC division during migration in mediating Laminin expression to orchestrate radial sorting and peripheral myelination *in vivo*.

## Introduction

Schwann cells are the myelinating glia that ensure efficient nerve impulse conduction along the nerves of the peripheral nervous system (PNS) (Boucanova & Chrast, 2020, Herbert & Monk, 2017, Jessen & Mirsky, 2005, Pereira, Lebrun-Julien et al., 2012, Raphael, Lyons et al., 2011, Sherman & Brophy, 2005, Woodhoo & Sommer, 2008). In order to myelinate, Schwann cells go through a series of developmental changes that include i) migration and division along peripheral axons ii) intensive proliferation prior to axonal ensheathment given that one Schwann cell myelinates one axonal segment and iii) substantial cytoskeletal rearrangements and changes in cell shape that allow a Schwann cell to radially sort an axon in a 1:1 ratio, a process termed radial sorting (Feltri, Poitelon et al., 2016, Monk, Feltri et al., 2015, Raphael & Talbot, 2011). The latter enables Schwann cells to extend their processes along a specific and unique abaxonal-adaxonal polarity axis in order to select and sort an axon to myelinate (Tricaud, 2017). It has been shown that *in vitro* Schwann cells first secrete and assemble their own basal lamina (BL) (Colognato & Tzvetanova, 2011, Eldridge, Bunge et al., 1989) that would later on regulate, through receptor interactions, different signals required for Schwann cell proliferation, survival and differentiation (Chen & Strickland, 2003, Court, Hewitt et al., 2009, Nodari, Previtali et al., 2008, Nodari, Zambroni et al., 2007, Yamada, Denzer et al., 1996, Yu, Feltri et al., 2005). Of particular interest are the extracellular matrix (ECM) proteins Laminins and Collagens that play essential roles in Schwann cell development (Chernousov, Yu et al., 2008).

However, several issues regarding the characteristics of timely Schwann cell division during migration and radial sorting, as well as the coupling of division with ECM proteins and myelination *in vivo* remain unresolved.

The mitotic spindle is a bipolar array of microtubules that mediates chromosome separation during cell division (Hara & Fukagawa, 2020, Petry, 2016). The organization and temporal assembly of the spindle are all important features that dictate the outcome of division (Lu & Johnston, 2013). One critical aspect of cell division is the temporal control of mitosis; this is highlighted by the existence of a mitotic spindle checkpoint that ensures accurate mitotic spindle organization prior to anaphase. It generates a wait signal until mitotic checkpoint proteins are removed from the kinetochore during metaphase by dynein/dynactin, clearing the way for accurate segregation of chromosomes into daughter cells (Lara-Gonzalez, Westhorpe et al., 2012). A great number of proteins is involved in spindle organization, one such major player is Sil that is ubiquitously expressed and specifically localizes to the poles of the mitotic spindle in metaphase cells (Campaner, Kaldis et al., 2005, Pfaff, Straub et al., 2007). The *sil* gene was originally cloned from leukemia-associated chromosomal translocation and is overexpressed in several tumor types such as melanoma and lung carcinomas (Aplan, Lombardi et al., 1990, Aplan, Lombardi et al., 1991). It is associated with autosomal recessive primary microcephaly (MCPH), a neurogenic mitotic disorder that results in significantly reduced brain size (Naveed, Kazmi et al., 2018, Zaqout, Morris-Rosendahl et al., 2017). Studies of zebrafish *cassiopeia* mutant *(csp^-/-^),* which has a nonsense mutation and a loss of function in zebrafish *sil,* show abnormalities in prometaphase progression within retinal neuroepithelium (Novorol, Burkhardt et al., 2013). Thus, *sil* represents an ideal candidate to study for a role in mitotic synchronization and division during Schwann cells’ development.

Here, we used live imaging, genetics and pharmacological tools in zebrafish to monitor Schwann cell behavior during its division and unraveled for the first time an essential role for early Schwann division during migration in peripheral myelination. We identified Sil as a critical regulator of Schwann cell radial sorting and myelination. Time-lapse imaging revealed an important role for *sil* in Schwann cell metaphase progression that delayed mitotic exit and caused a significant decrease of peripheral myelin markers, *krox20* and *mbp* in *csp^-/-^* embryos. This coincided with a complete loss of axonal ensheathment as revealed by transmission electron microscopy (TEM). Treating *csp^-/-^* with forskolin that binds to adenyl cyclase and elevates the levels of cAMP, restored radial sorting, myelin-associated genes expression and axonal wrapping. Moreover, *csp^-/-^* mutants failed to express Laminin within the posterior lateral line nerve (PLLn) and forcing Laminin2 expression was sufficient to fully restore peripheral myelination. These data unraveled an essential role for the timely process of mitotic exit within Schwann cells, specifically during migration, in radial sorting and myelination via Laminin2/cAMP dependent pathway. We provide evidence that Schwann cells have a limited window of time in which they have to divide during their migration in order to initiate radial sorting and myelination.

## Results

### Schwann cells show a regular pattern of division during radial sorting along the PLLn

In order to monitor Schwann cell division, we took advantage of the *Tg(foxd3:gfp)* (Gilmour, Maischein et al., 2002) and first imaged Schwann cells along the PLLn between 48 and 60 hours post-fertilization (hpf). During this time, Schwann cells divide intensively and start to radially sort axons that precedes axonal ensheathment (Figure S1) (Lyons, Pogoda et al., 2005, Raphael et al., 2011). Schwann cells were generally elongated along the axons until they went through mitotic rounding that led to division and then re-elongated again (Figure 1A). Cells divided along the Anterior-Posterior (AP) axis of the embryo and it took an average of 9.30±0.61 min for mitosis to complete (Figure 1B and Movie S1). To assess the importance of this pattern of division in Schwann cell myelination, we decided to monitor their behavior in *cassiopeia* mutant *(csp^-/-^)*. Sil is a ubiquitously expressed protein that specifically localizes to the mitotic spindle in metaphase cells and plays a critical role in its organization (Novorol et al., 2013, Pfaff et al., 2007). Schwann cells’ mitosis in *csp^-/-^* took much longer to complete with an average of 90.10±14.13 min in comparison to 9.30±0.61 min in controls (Figure 1A,B and Movie S2). However, re-elongation of Schwann cells occurred normally after division in both mutants and controls (Figure 1A).

**Figure 1.**
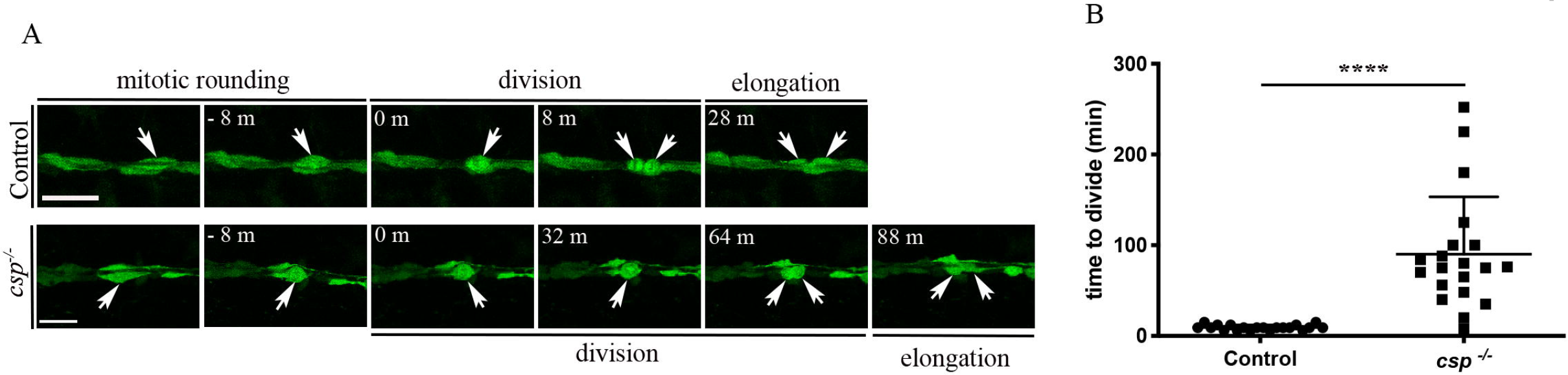
Schwann cells show a regular pattern of division during radial sorting (A) Still images of time-lapse imaging in *Tg(foxd3:gfp)* control and *Tg(foxd3:gfp)/csp^-/-^* embryos. Arrows indicate Schwann cells along the PLLn at different timepoints prior to and after division. Scale bars = 25μm. (B) Quantification of the time required for control (20 cells, n= 6 embryos) and *csp^-/-^* (20 cells, n= 5 embryos) Schwann cells to successfully complete mitotic division. (****, p ≤ 0.0001). m or min, minutes.

This result suggests that Sil deficiency leads to a significant delay in Schwann cell mitosis.

### Sil is essential for Schwann cell radial sorting and myelination

Having identified a delay in progression through mitosis in *csp^-/-^* Schwann cells, we wondered whether this defect could have any impact on the ability of Schwann cells to ensheath axons. Transmitted Electron Microscopy (TEM) analysis showed a dramatic decrease in the number of myelinated axons in *csp^-/-^* in comparison to controls. We observed an average of 5.6±0.55 myelinated axons per nerve in controls in comparison to 0 myelinated axon per nerve in *csp^-/-^* at 72 hpf (or 3 days post-fertilization (dpf)) (Figure 2A,B,E). In addition, we observed a significant decrease in the total number of axons in these mutants with an average of 55±2.85 axons in controls and 28.63±2.67 in *csp^-/-^* PLLn (Figure 2F). In order to test whether this defect is maintained at later stages, we analyzed axonal wrapping at 4 dpf. The myelination defect in *csp^-/-^* embryos detected at 3 dpf was also observed at 4 dpf (Figure 2C-F). This analysis could not be extended to later stages since *csp^-/-^* embryos died between 4 and 5 dpf.

**Figure 2.**
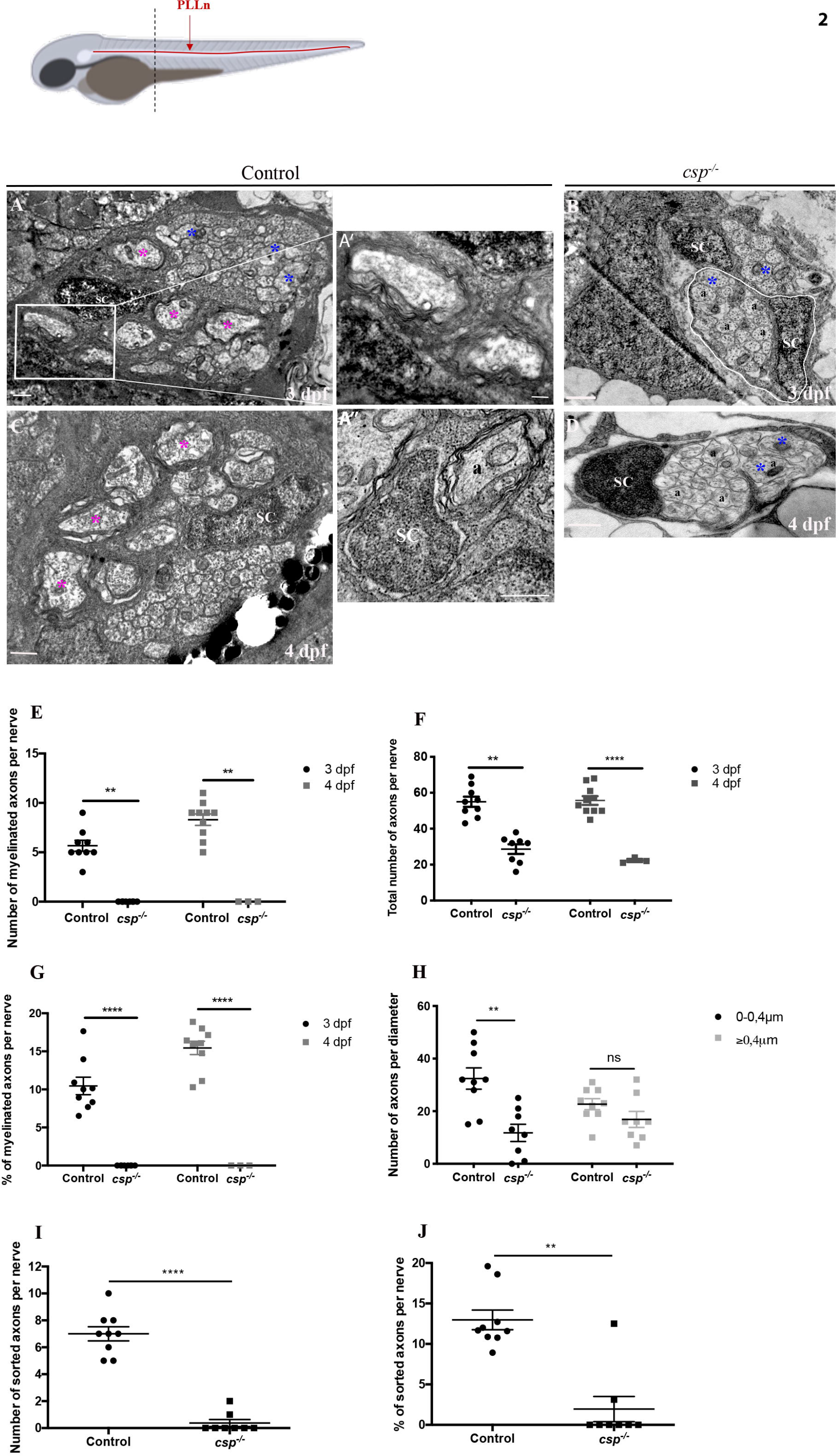
Sil is essential for radial sorting and myelination by Schwann cells Schematic illustration of a zebrafish larvae. PLLn is shown in red. Dotted line represents the anterior-posterior position of cross section analysis in TEM. PLLn, posterior lateral line nerve. TEM of a cross section of the PLLn at 3 dpf in control (A) and *csp^-/-^* embryo (B) and at 4 dpf in control (C) and *csp^-/-^* (D). Magenta asterisks highlight some large caliber myelinated axons (some shown at higher magnification in A’, scale bar = 0.2 μm) and blue asterisks show some large caliber non-myelinated axons. Scale bars = 0.5 μm. SC, Schwann cell. (A’’) Example of a 1:1 association between a Schwann cell and an axon, i.e. a radially sorted axon in control embryo at 3 dpf. Scale bar = 0.5 μm. a, axon. Axons remained bundled in *csp^-/-^* whereby one Schwann cell is associated with a bundle of axons, delimitated in white in B. (E) Quantification of the number of myelinated axons per nerve at 3 dpf in controls (9 nerves, n= 5 embryos) and *csp^-/-^* (8 nerves, n= 6 embryos) and at 4 dpf in controls (average of 8.3±0.57 myelinated axons; 10 nerves, n= 6 embryos) and *csp^-/-^* (0 myelinated axon; 3 nerves, n= 3 embryos) (**, p= 0.0035 at 3 dpf; **, p= 0.005 at 4 dpf). (F) Quantification of the total number of axons per nerve at 3 dpf in controls and *csp^-/-^* and at 4 dpf in controls (54±3.16 axons) and *csp^-/-^* (23±1 axons) (**, p= 0.003; ****, p≤0.0001). (G) Quantification of the percentage of myelinated axons relative to the total number of axons per nerve at 3 dpf in controls and *csp^-/-^* and at 4 dpf in controls (15.44±0.87%) and *csp^-/-^* (0%) (****, p≤0.0001; ****, p≤0.0001). (H) Quantification of the number of axons relative to their diameter at 3 dpf in controls and *csp^-/-^* (**, p= 0.0013; ns, p= 0.14). (I) Quantification of the number of sorted axons per nerve at 3 dpf in controls (average of 7.00±0.52) and *csp^-/-^* (0.37±0.26) (****, p≤0.0001). (J) Quantification of the percentage of sorted axons relative to the total number of axons at 3 dpf in controls (12.97±1.21) and *csp^-/-^* (1.95±1.55) (**, p=0.0019).

Given the drastic decrease in total number of axons in these mutants, we therefore calculated the ratio of myelinated axons relative to the total number of axons in the PLLn. While we counted an average of 10.46±1.14% of myelinated axons in controls, we observed 0% in *csp^-/-^* at 3 and 4 dpf (Figure 2G). This myelination defect was not the result of a decrease in the number of large caliber axons that are supposed to be myelinated (Figure 2H). Finally, the myelination defect coincided with a significant decrease in the number and the percentage of radially sorted axons per nerve (Figure 2I,J).

These results show that *sil* is required for Schwann cell radial sorting and myelination.

### Sil is required within Schwann cells for axonal wrapping and normal Mbp expression and regulates the number of neurons within the PLLg

*Sil* is ubiquitously expressed but highly enriched in neural cells and *csp^-/-^* embryos show major neuronal defects such as an increase in the percentage of mitotically arrested neuroepithelial cells in the retina that coincides with an increase in apoptosis (Novorol et al., 2013). To test whether the myelination defect observed here is specific to Schwann cell or secondary to neuronal defects, we first examined several aspects of posterior lateral line ganglia (PLLg) and PLLn development in these mutants. *Csp^-/-^* embryos presented no major defects in PLLn growth and Schwann cell migration at 48 hpf along the PLLn (Figure S2A). A few *csp^-/-^* embryos (< 30%, n=7/24) showed an axonal growth arrest but posterior to the yolk extension and Schwann cells were found at the tip end of these axons. We next analyzed axonal transport along the PLLn using *mito:gfp* construct that labels mitochondria and we monitored their behavior by time-lapse imaging. We observed no significant difference in the average speed of mitochondria along the PLLn with an average of 1.92±0.14 μm/s and 1.85±0.06 μm/s in controls and *csp^-/-^* respectively (Figure S2B,C and Movies S3, S4). We next looked at the PLLg and counted the number of neurons within the ganglia using the neuronal marker HuC. We observed a significant decrease in the number of neurons in *csp^-/-^* embryos at 48 and 72 hpf in comparison to controls (Figure S2D,E) suggesting a role for *sil* in the development of PLLg that coincides with the reduction in the number of axons in the PLLn.

These results suggest a role for *sil* in the development of PLLg but not in axonal growth or transport.

To directly test whether *sil* has an autonomous function in Schwann cell myelination, we forced the expression of *sil* specifically in Schwann cells under the control of the *sox10* promoter. *Csp^-/-^* embryos were injected with pTol2-*sox10:sil-P2A-mcherry-CaaX* and tol2 transposase mRNA at 1 cell stage and selected at 3 dpf for *mcherry* expression in Schwann cells to be later fixed for TEM analysis. By forcing the expression of *sil* specifically in Schwann cells, we significantly increased the number of myelinated axons in *csp^-/-^* embryos at 3 dpf (Figure 3A-B’’). The percentage of myelinated axons per nerve in *csp^-/-^* injected embryos was similar to the one observed in controls (Figure 3C).

**Figure 3.**
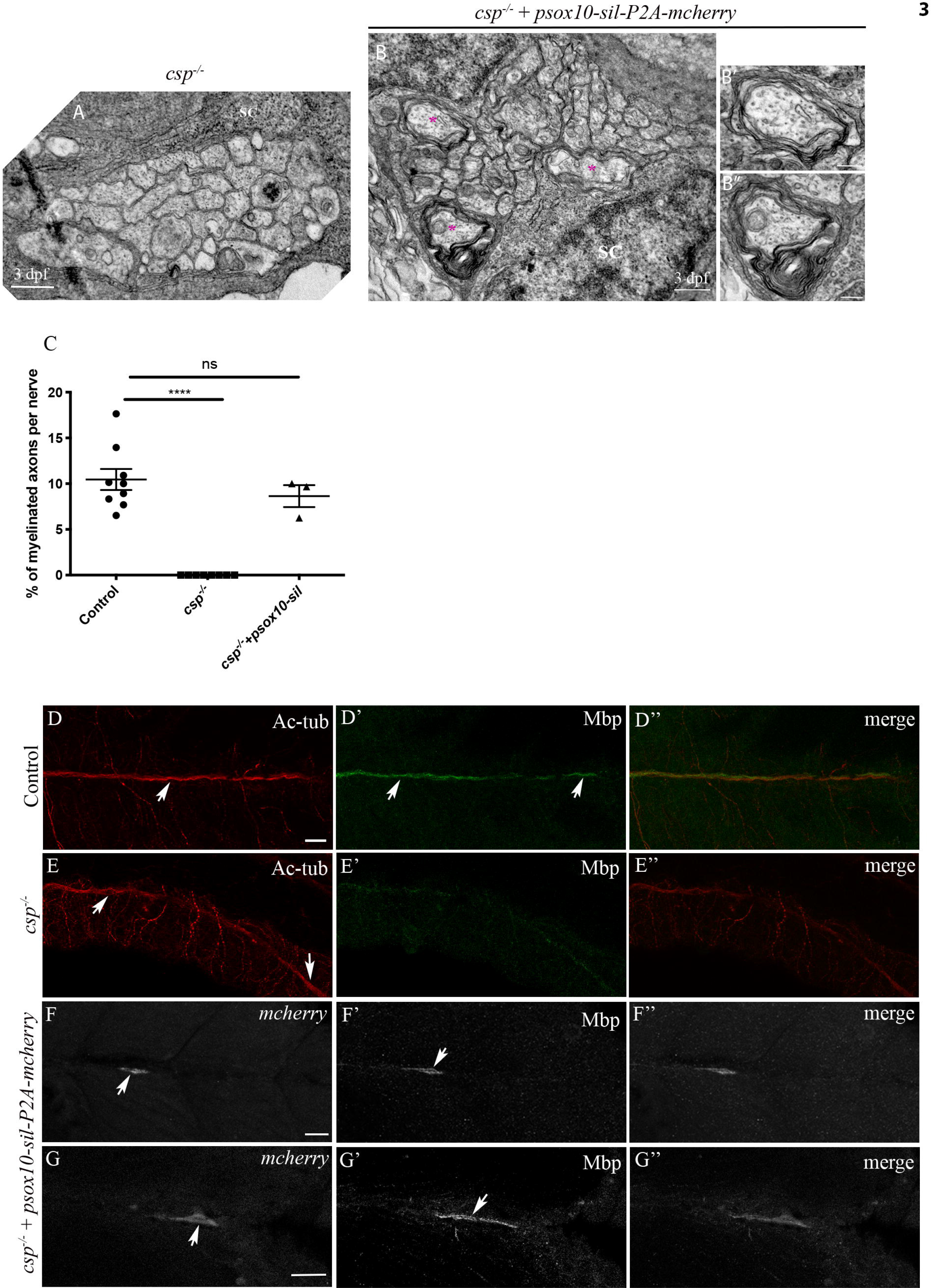
Sil is required within Schwann cells for axonal wrapping TEM of a cross section of the PLLn at 3 dpf in *csp^-/-^* (A) and *csp^-/-^* embryos injected with *psox10-sil-P2A-mcherry* (B). Magenta asterisks represent some large caliber myelinated axons (some shown at higher magnification in B’, B’’, scale bars = 0.2 μm). Scale bars = 0.5 μm. (C) Quantification of the percentage of myelinated axons relative to the total number of axons in control (average of 10.46±1.14%, 9 nerves, n= 5 embryos), *csp^-/-^* (0%, 8 nerves, n= 6 embryos) and *csp^-/-^* injected with *psox10-sil-P2A-mcherry* (average of 8.64±1.20%, 3 nerves, n= 3 embryos) (****, p≤0.0001; ns, p= 0.4871). Acetylated tubulin (Ac-tub) and Mbp immuno-labeling in control (D-D’’), *csp^-/-^* (E-E’’) and *csp^-/-^+psox10-sil-P2A-mcherry* (F-G’’) embryos at 3 dpf. Arrows indicate the PLLn in D and E, the myelin sheaths in D’, mcherry+ Schwann cells along the PLLn in F and G and the corresponding Mbp expression in F’ and G’. D’’, E’’, F’’ and G’’ are the corresponding merge images of D and D’, E and E’, F and F’, G and G’ respectively. Scale bars = 20 μm

We also processed control and *csp^-/-^* larvae for immunohistochemistry to detect Mbp at 3 dpf. We observed a sharp decrease in Mbp expression in *csp^-/-^* embryos in comparison to controls (Figure 3D-E’’). We then injected *csp^-/-^* embryos with pTol2-*sox10:sil-P2A-mcherry-CaaX* and tol2 transposase mRNA at 1 cell stage and selected embryos for mosaic mcherry+ clonal analysis. These embryos were then processed for immunohistochemistry to detect Mbp. We analyzed 4 mutant larvae that had *sox10*:mcherry^+^ Schwann cells along mutant PLLn; the mcherry+ clones appeared to express normal levels of Mbp (Figure 3F-G’’).

Altogether, these results suggest a specific requirement for *sil* within Schwann cells to initiate axonal wrapping and Mbp expression.

### Schwann cells in *csp^-/-^* show delays in mitotic progression but exit mitosis with no significant increase in apoptosis

Given that *sil* deficiency leads to a decrease in the number of neurons with significant apoptosis in the retina, it is possible that the myelination defects observed in the PLLn of *csp^- /-^* embryos could be related to a defect in the number of Schwann cells and/or their survival.

To address this question, we first labeled Schwann cells using the M phase marker PH3 in *Tg(foxd3:gfp)* at 48 and 72 hpf. As seen in the retina (Novorol et al., 2013), we observed a significant increase in the fraction of proliferating PH3+/Schwann cells at 48 hpf. This coincided with a decrease in the total number of Schwann cells and an increase in the ratio of PH3+ Schwann cells relative to the total number of Schwann cells counted within a region of 8 somites (Figure 4A-D). However, results were different at 72 hpf, as the total number of Schwann cells, the number of PH3+/Schwann cells and the ratio of PH3+/Schwann cells relative to the total number of Schwann cells showed no significant difference between controls and *csp^-/-^* embryos (Figure 4A-D). Moreover, time-lapse analysis using the nuclear marker *h2b:gfp* pointed to a delay in prometaphase progression (Figure 4E and Movies S5, S6). However, in contrast to the neuronal phenotype in retina, most of Schwann cells exited mitosis and we detected no apoptotic Schwann cells in these mutants along the PLLn (Figure 4F,G). We also observed a significant increase in the number of acridine orange (AO)+ cells within the spinal cord that is comparable to the defect observed in the zebrafish retina suggesting a role for *sil* in cell survival outside the brain (Figure 4F,H).

**Figure 4.**
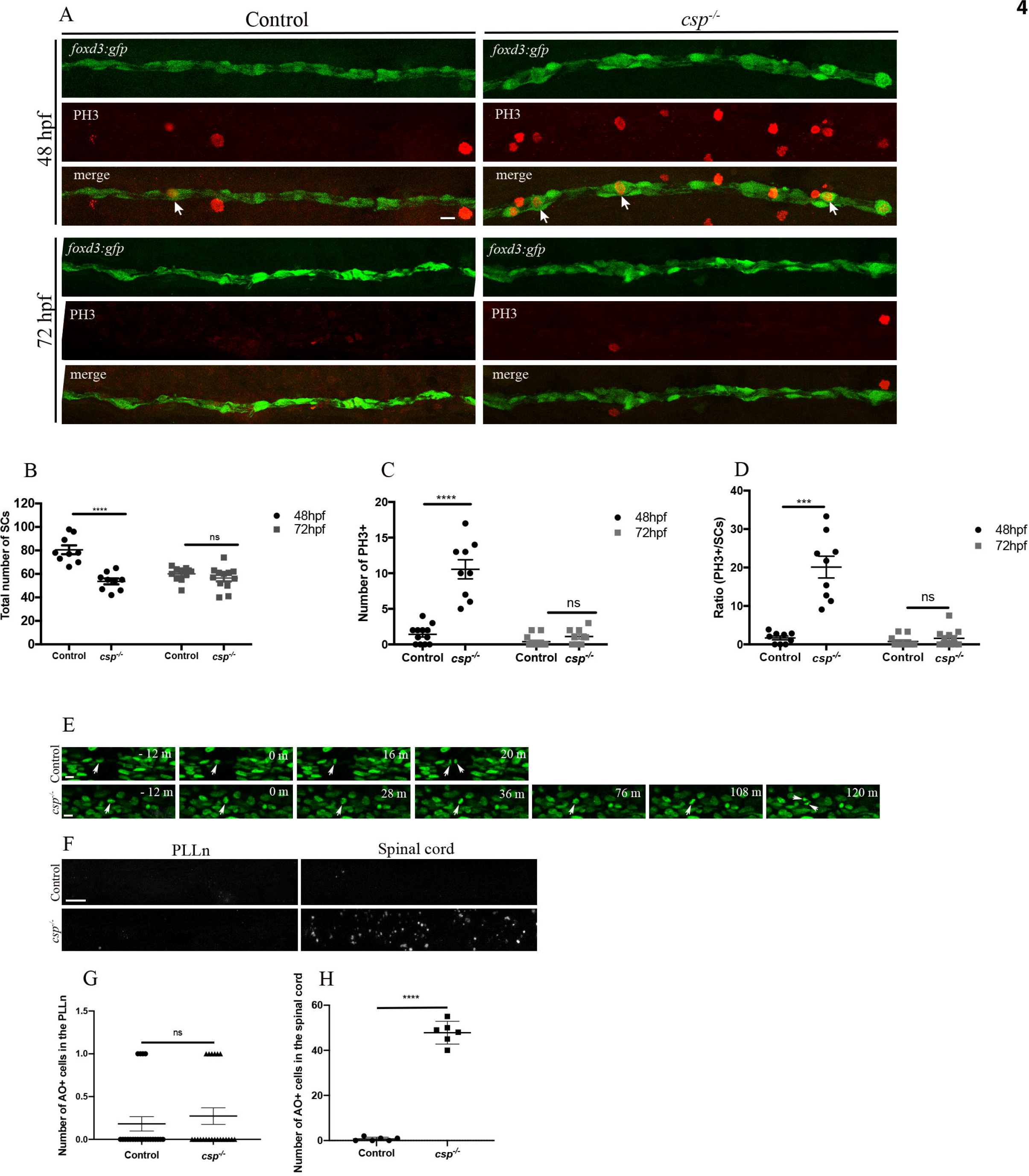
Schwann cells in *csp^-/-^* show delays in mitotic progression but exit mitosis with no significant increase in apoptosis (A) PH3 immuno-labeling in *Tg(foxd3:gfp)* and *Tg(foxd3:gfp)/csp^-/-^* at 48 hpf and 72 hpf. Arrows indicate Schwann cells that are GFP and PH3 positives. Scale bar = 25 μm. (B) Quantification of the number of Schwann cells within a defined region of the PLLn at 48 and 72 hpf in control (average of 80.56±3.76 cells at 48 hpf, n= 9 embryos; average of 60±1.78 cells at 72 hpf, n= 11 embryos) and *csp^-/-^* (average of 53.57±2.50 at 48 hpf, n= 9 embryos; average of 56.42±2.75 cells at 72 hpf, n= 12 embryos) (****, p≤ 0.0001; ns, p= 0.2897). (C) Quantification of the number of PH3^+^/Schwann cells within a defined region of the PLLn at 48 and 72 hpf in control (average of 1.41±0.37, n=12 embryos at 48 hpf; average of 0.38±0.23, n= 13 embryos at 72hpf) and *csp^-/-^* (average of 10.56±1.34, n=9 embryos at 48 hpf; average of 1.11±0.35, n= 9 embryos at 72 hpf) (****, p≤ 0.0001; ns, p= 0.064). (D) Quantification of the ratio of PH3^+^ Schwann cells relative to the total number of Schwann cells within a defined region of the PLLn at 48 and 72 hpf in control (average of 1.68±0.48, n= 9 embryos at 48 hpf; average of 0.75±0.41, n= 11 embryos at 72hpf) and *csp^-/-^* (average of 20.13±2.82, n= 9 embryos at 48 hpf; average of 1.59±0.64, n= 12 embryos at 72 hpf) (***, p= 0.0002; ns, p= 0.3636). (E) Still images of time-lapse imaging in control (average of 16.67±1.22 min, 6 nuclei, n= 3 embryos) and *csp^-/-^* (average of 98.67±4.55 min, 6 nuclei, n= 3 embryos) embryos injected with *h2b:gfp.* Arrows indicate Schwann cells nuclei from the beginning of M phase (time 0) when mitotic rounding takes place until the two nuclei split. Scale bars = 10 μm. m, minutes. (F) Acridine orange (AO) staining at 50 hpf in control and *csp^-/-^* embryos within a defined region of the PLLn and spinal cord. Scale bar = 25 μm. (G) Quantification of the number of AO positive cells in control (average of 0.18±0.08, n= 11 embryos) and *csp^-/-^* (average of 0.27±0.09, n= 11 embryos) within a defined region of the PLLn (ns, p= 0.72). (H) Quantification of the number of AO positive cells in control (average of 0.66±0.33, n= 6 embryos) and *csp^-/-^* (average of 47.83±2.05, n= 6 embryos) within a defined region of the spinal cord (****, p≤0.0001).

These data suggest that *sil* is required in Schwann cells for the temporal control of mitotic exit, nevertheless, Schwann cells devoid of *sil* do exit mitosis following a significant delay and are not apoptotic.

### Cell division during radial sorting is not required *per se* for Schwann cell myelination

Since Schwann cells in *csp^-/-^* embryos show a delay in their mitotic progression, it is possible that there is a window of time during radial sorting in which Schwann cell have to divide in order to myelinate. Moreover, it has been shown that blocking cell division during radial sorting can alter their ability to radially sort axons and myelinate (Raphael et al., 2011). To test this hypothesis, we treated embryos with aphidicolin that blocks cell division *in vivo* in zebrafish (Lyons et al., 2005, Raphael et al., 2011). First, we incubated embryos in aphidicolin between 45 and 54 hpf (see Materials and Methods for aphidicolin treatment/efficacy) to temporarily inhibit cell division during the initial phase of radial sorting. The medium was then washed and embryos were allowed to develop until 72 hpf (or 3 dpf) to be fixed for TEM. Another group of embryos was incubated from 45 hpf until 72 hpf to block Schwann cell division during the whole period of radial sorting. TEM analysis showed a significant decrease in the number of myelinated axons in both groups of treated embryos but also, as expected, in the total number of axons within the PLLn (Figure 5A-E). However, some large caliber axons were myelinated. Most importantly, the ratios of myelinated axons as well as the ratios of axons per diameter relative to the total number of axons showed no significant difference between the different groups (Figure 5F-H).

**Figure 5.**
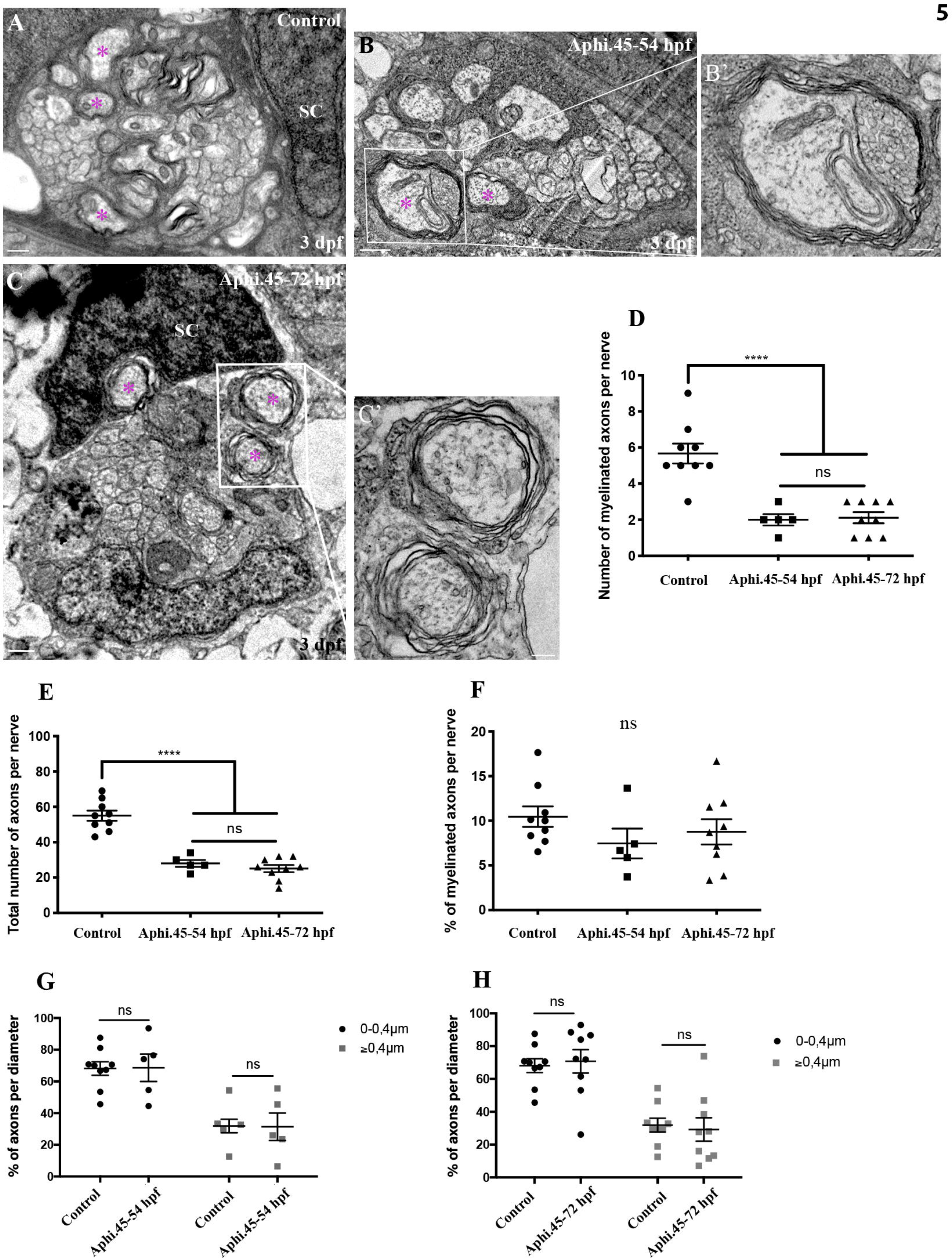
Schwann cell division during radial sorting is not required *per se* for myelination by Schwann cells TEM of a cross section of the PLLn at 3 dpf in control (A), embryo treated with aphidicolin between 45 and 54 hpf (B) and embryo treated with aphidicolin between 45 and 72 hpf (C). Magenta asterisks represent some large caliber myelinated axons, (some shown at higher magnification in B’ and C’, scale bars = 0.2 μ). SC, Schwann cell. Scale bars = 0.5 μm. (D) Quantification of the number of myelinated axons per nerve at 3 dpf in controls (average of 5.7±0.56, 9 nerves, n= 6 embryos), embryos treated with aphidicolin between 45 and 54 hpf (average of 2.11±0.31, 5 nerves, n= 5 embryos), and embryos treated with aphidicolin between 45 and 72 hpf (average of 2.00±0.32, 9 nerves, n= 5 embryos) (****, p≤0.0001; ns, p= 0.9790). (E) Quantification of the total number of axons per nerve at 3 dpf in controls (average of 57±2.9), embryos treated with aphidicolin between 45 and 54 hpf (average of 25.11±2.04) and embryos treated with aphidicolin between 45 and 72 hpf (average of 28.00±1.98) (****, p≤0.0001; ns, p= 0.6574). (F) Quantification of the percentage of myelinated axons relative to the total number of axons per nerve at 3 dpf in controls (average of 10.5±1.2), embryos treated with aphidicolin between 45 and 54 hpf (average of 8.76±1.42) and embryos treated with aphidicolin between 45 and 72 hpf (average of 7.46±1.66). ns, p> 0.05 in all cases. (G) Quantification of the percentage of axons according to their diameter relative to the total number of axons per nerve at 3 dpf in controls (average of 68.12 for 0-0.4 μm; average of 31.88 for >0.4 μm) and aphidicolin treated embryos between 45 and 54 hpf (average of 68.62 for 0-0.4 μ average of 31.38 for >0.4 μm). ns, p=0.9607. (H) Quantification of the percentage of axons according to their diameter relative to the total number of axons per nerve at 3 dpf in controls (average of 68.12 for 0-0.4 μm; average of 31.88 for >0.4 μm) and aphidicolin treated embryos between 45 and 72 hpf (average of 70.76 for 0-0.4 μm; average of 29.24 for >0.4 μm). ns, p=0.7545.

This result suggests that cell division during radial sorting is not required *per se* for peripheral myelination.

### Sil is required for mitotic temporal exit during Schwann cell migration

Given that cell division during radial sorting is not required *per se* for Schwann cell myelination, we therefore tested whether *sil* is also implicated in early Schwann cell division during migration. For this, we monitored Schwann cell behavior during axonal growth when Schwann cells migrate and divide along the axons of the PLLn. Schwann cells were generally elongated during migration too until they went through mitotic rounding, divided and then re-elongated (Figure 6A, Movie S7). Schwann cells divided along the AP axis of the embryo and it took an average of 9.81±0.63 min for them to complete mitosis, showing a very similar behavior to the one observed during radial sorting. However, Schwann cells struggled to exit mitosis in *csp^-/-^* with an average of 87.20±4,72 min for mitosis to be completed (Figure 6A,B, Movie S8).

**Figure 6.**
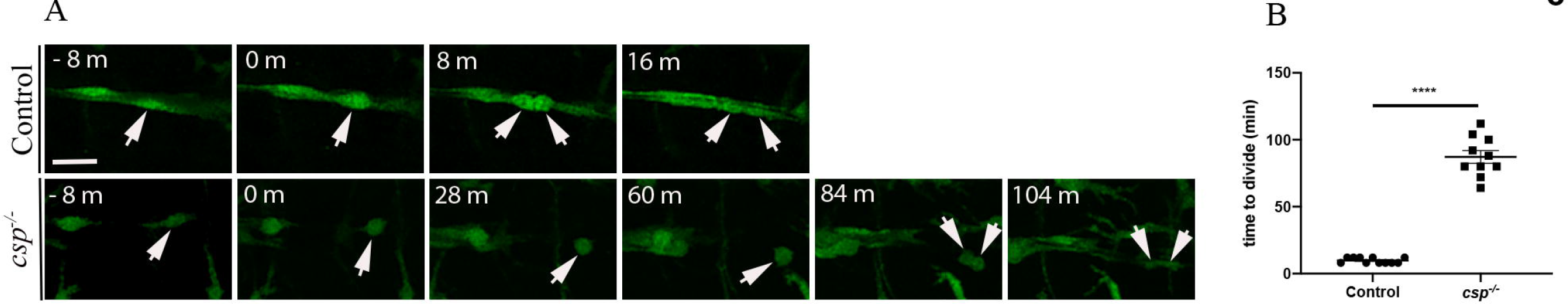
*Sil* is required for Schwann cell temporal mitotic exit during migration (A) Still images of time-lapse imaging in *Tg(foxd3:gfp)* control and *Tg(foxd3:gfp)/csp^-/-^* embryos at around 30 hpf. Arrows indicate Schwann cells along the PLLn at different timepoints prior to and after division. Scale bar = 20 μm. (B) Quantification of the time required for control (11 cells, n= 3 embryos) and *csp^-/-^* (10 cells, n= 3 embryos) Schwann cells to successfully complete mitotic division during migration. (****, p ≤ 0.0001). m or min, minutes.

This result shows that *sil* is also required for the temporal control of mitosis in Schwann cell during migration similar to what we observed during radial sorting.

### Early Schwann cell division during migration is essential for peripheral myelination

Since *sil* is also required for the temporal control of Schwann cell mitosis during migration, it is possible that the myelination defects observed in *csp^-/-^* result from a defective division during these earlier events. It has been shown that Schwann cell division during migration is not required for migration *per se* (Lyons et al., 2005), however whether it is required for myelination has not been tested yet. We therefore treated embryos with aphidicolin to block cell division specifically during Schwann cell migration (between 22 and 40 hpf) and before radial sorting. The medium was then washed and embryos were allowed to develop until 3 dpf to be fixed for TEM. As expected, there was a sharp decrease in the total number of axons per nerve, however, we counted an average of only 0.125 myelinated axon in aphidicolin treated embryos at 3 dpf (Figure 7A-E). To test whether Schwann cells were able to divide again during radial sorting once the medium is washed, we analyzed their division and numbers during radial sorting, here at 52 hpf. We observed a significant decrease in the number of Schwann cells in treated embryos at 52 hpf but the number of Schwann cells that are PH3+ and the ratio of PH3+ Schwann cells relative to the total number of Schwann cells were similar between controls and treated embryos (Figure 7F-H). This result shows that Schwann cells were able to divide again during radial sorting following the temporal cell division block during migration.

**Figure 7.**
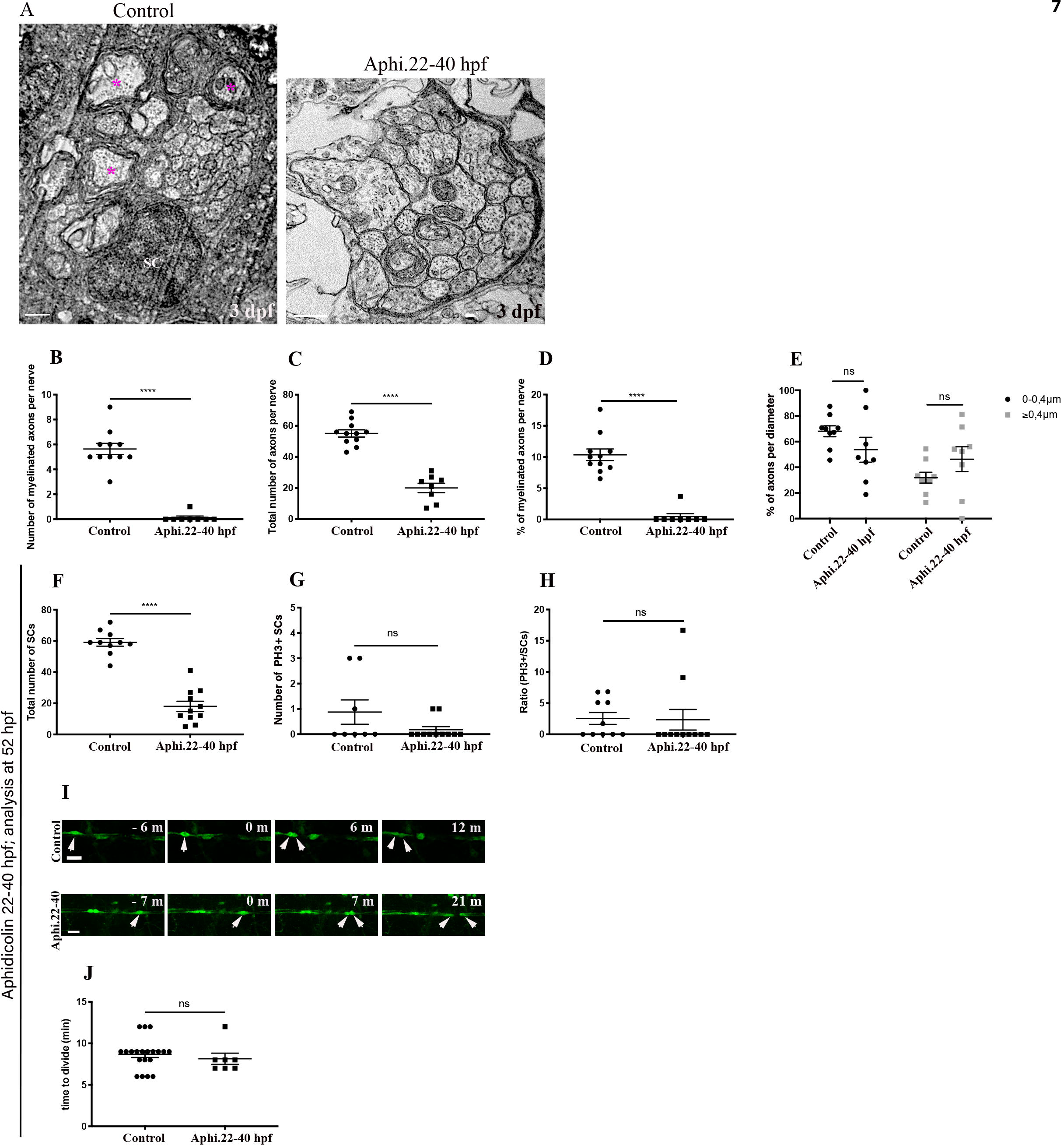
Early Schwann cell division during migration is essential for myelination (A) TEM of a cross section of the PLLn at 3 dpf in control and embryo treated with aphidicolin between 22 and 40 hpf. Magenta asterisks represent some large caliber myelinated axons. SC, Schwann cell. Scale bars = 0.5 μm. (B) Quantification of the number of myelinated axons per nerve at 3 dpf in controls (average of 5.63±0.45, 11 nerves, n= 7 embryos) and embryos treated with aphidicolin between 22 and 40 hpf (average of 0.125±0.125, 8 nerves, n= 6 embryos) (****, p≤0.0001). (C) Quantification of the total number of axons per nerve at 3 dpf in controls (average of 55.09±2.32) and embryos treated with aphidicolin between 22 and 40 hpf (average of 20±3.04) (****, p≤0.0001). (D) Quantification of the percentage of myelinated axons relative to the total number of axons per nerve at 3 dpf in controls (average of 10.36±0.93) and embryos treated with aphidicolin between 22 and 40 hpf (average of 0.46±0.46) (****, p≤0.0001). (E) Quantification of the percentage of axons according to their diameter relative to the total number of axons per nerve at 3 dpf in controls (average of 68.12 for 0-0.4 μm; average of 31.88 for >0.4 μm) and aphidicolin treated embryos between 22 and 40 hpf (average of 53.69 m; average of 46.31 for >0.4 μm). ns, p=0.2045. (F) Quantification of the number of Schwann cells within a defined region of the PLLn at 52 hpf in control (average of 59.10±2.44 cells, n= 10 embryos) and embryos treated with aphidicolin between 22 and 40 hpf (average of 18±3.27 cells, n= 11 embryos) (****, p≤ 0.0001). (G) Quantification of the number of PH3^+^/Schwann cells within a defined region of the PLLn at 52 hpf in control (average of 0.87±0.47, n=8 embryos) and embryos treated with aphidicolin between 22 and 40 hpf (average of 0.18±0.12, n= 11 embryos) (ns, p= 0.3265). (H) Quantification of the ratio of PH3^+^ Schwann cells relative to the total number of Schwann cells within a defined region of the PLLn at 52 hpf in control (average of 2.54±0.95, n= 10 embryos) and embryos treated with aphidicolin between 22 and 40 hpf (average of 2.34±1.65, n= 11 embryos) (ns, p= 0.3274). (I) Still images of time-lapse imaging in *Tg(foxd3:gfp)* control and *Tg(foxd3:gfp)* treated with aphidicolin between 22 and 40 hpf embryos at around 52 hpf. Arrows indicate Schwann cells along the PLLn at different timepoints prior to and after division. Scale bars = 20 μm. (J) Quantification of the time required for control (average of 8.70±0.41 min, 20 cells, n= 5 embryos) and embryos treated with aphidicolin between 22 and 40 hpf (average of 8.14±0.67 min, 7 cells, n= 4 embryos) Schwann cells to successfully complete mitotic division between 48 and 60 hpf. (ns, p= 0.1796).

We also monitored Schwann cell division during radial sorting using live imaging in controls and treated embryos. Our results showed a normal pattern of division during radial sorting in treated embryos similar to controls (Figure 7I,J, Movies S9,10).

Altogether, these results point to an essential role for Schwann cell division during migration in myelination.

### *Sil* is required to initiate radial sorting, myelin gene expression and axonal wrapping by Schwann cells via cAMP pathway

M phase is characterized by chromatin condensation and temporal ejection of transcription factors (Ma, Kanakousaki et al., 2015). Since *sil* is implicated in spindle checkpoint and mitotic progression during M phase, it is possible that the delayed mitotic exit observed in *csp^-/-^* embryos could result in prolonged ejection of transcription factors required for myelination in Schwann cells. Alternatively, it is possible that transcriptional activity is effective after mitosis but Schwann cells in *csp^-/-^* embryos present defective upstream signaling activity. To distinguish between these possibilities, we first analyzed the expression of *krox20* and *mbp* in controls and *csp^-/-^* embryos at 52 hpf and 3 dpf respectively. *csp^-/-^* embryos showed a sharp decrease in the expression of peripheral myelin genes (Figure 8B,E) in comparison to controls (Figure 8A,D) confirming the role of *sil* in this process. This coincided, as shown before, with a total lack of radial sorting and axonal ensheathment (Figure 8H). Second, we decided to treat embryos with forskolin that interacts with adenyl cyclase and elevates the levels of cAMP. It has been shown that elevation of cAMP *in vivo* in zebrafish is able to re-initiate the defective expression of myelin transcription factors in *gpr126* mutant (Monk, Naylor et al., 2009). Indeed, when *csp^-/-^* embryos were treated with forskolin during radial sorting, the expression of *krox20* and *mbp* was restored (Figure 8C,F) as well as the ratio of myelinated axons relative to the total number of axons (Figure 8I,J).

**Figure 8.**
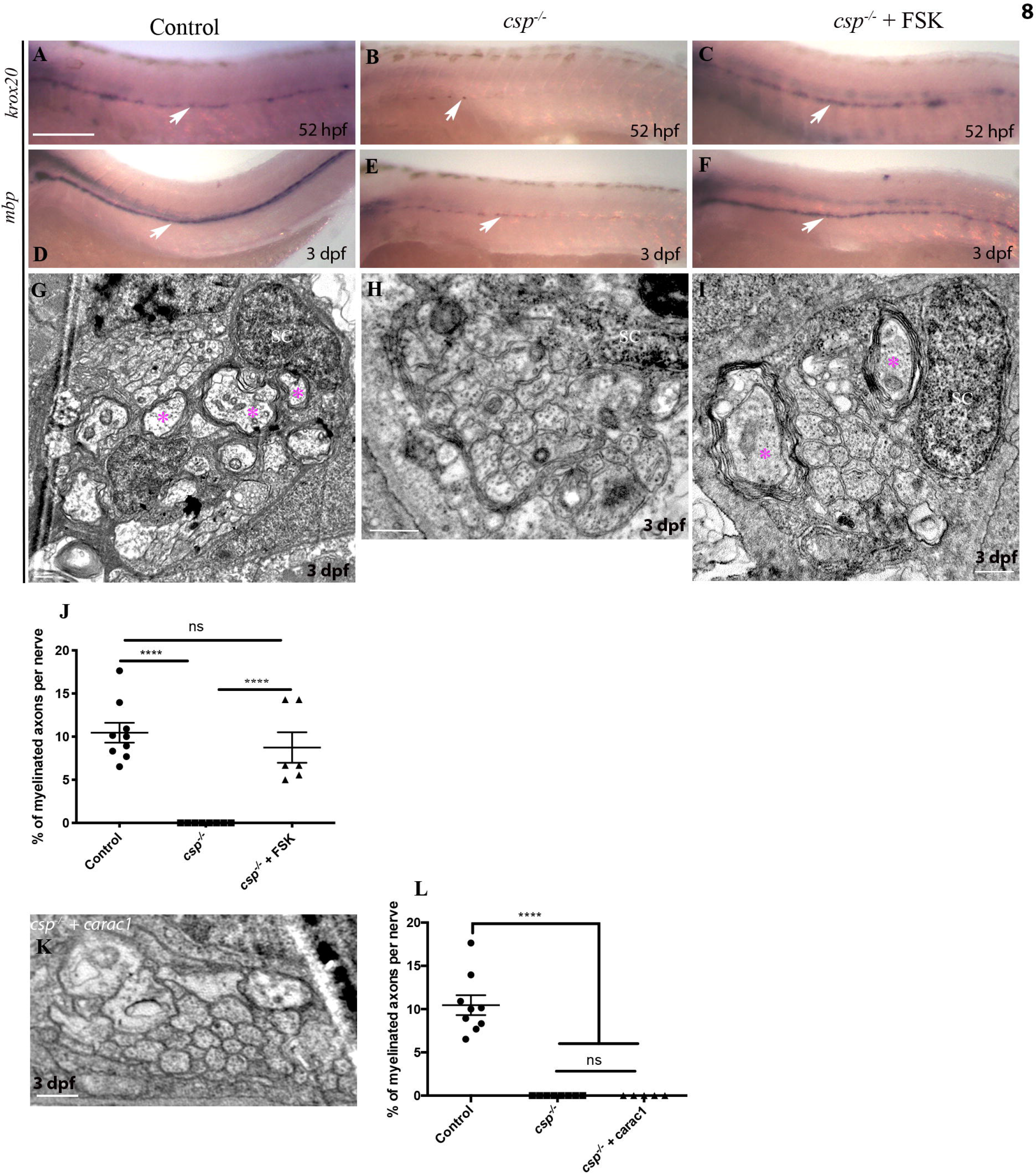
*Sil* is required to initiate myelin gene expression and axonal wrapping by Schwann cells via cAMP dependent pathway Lateral views of *krox20* expression at 52 hpf revealed by in situ hybridization along the PLLn in control (A) showing a robust expression (n=29), *csp^-/-^* embryo (B) showing a sharp decrease in *krox20* expression (n=22) and in *csp^-/-^* treated with forskolin (FSK) (C) showing a robust expression similar to controls (n=30). Arrows indicate Schwann cells expressing *krox20* along the PLLn. Lateral views of *mbp* expression at 3 dpf revealed by in situ hybridization along the PLLn in control (D) showing a robust expression (n=31), *csp^-/-^* embryo (E) showing a sharp decrease in *mbp* expression (n=24) and in *csp^-/-^* treated with forskolin (F) showing a robust expression similar to controls (n=28). Arrows indicate Schwann cells expressing *mbp* along the PLLn. Scale bar = 200 μm. TEM of a cross section of the PLLn at 3 dpf in control (G), *csp^-/-^* (H) and *csp^-/-^* treated with forskolin between 45 and 54 hpf (I). Magenta asterisks represent some large caliber myelinated axons. SC, Schwann cell. Scale bars = 0.5 μm. (J) Quantification of the percentage of myelinated axons relative to the total number of axons per nerve at 3 dpf in controls (average of 10.6±1.17), *csp^-/-^* (average of 0) and *csp^-/-^* treated with forskolin (average of 8.74±1.77) (****, p≤0.0001; ns, p= 0.4846; ****, p≤0.0001). (K) TEM of a cross section of the PLLn at 3 dpf in *csp^-/-^* embryo injected with a constitutive active form of Rac1 (*carac1*). Scale bar = 0.5 μm. (L) Quantification of the percentage of myelinated axons relative to the total number of axons per nerve at 3 dpf in controls (average of 10.5±1.13), *csp^-/-^* (average of 0) and *csp^-/-^* injected with *carac1* (average of 0) (****, p≤0.0001, ns, p≥ 0.999).

This result suggests that the myelin defects observed in *csp^-/-^* embryos result from a defective cAMP signaling pathway and not from a prolonged ejection of transcription factors following a particularly lengthy M phase.

### Forcing Rac1 activity does not rescue radial sorting and myelination defects in *csp^-/-^*

The radial sorting defect and lack of axonal ensheathment by Schwann cell observed in this mutant might also be linked to a defective cytoskeletal re-arrangement mediated by Rac1. To test this hypothesis, we forced the expression of Rac1, a small GTPase protein that interacts with Schwann cell cytoskeleton (Nodari et al., 2007) and is able to restore peripheral radial sorting defects in zebrafish (Boueid, Mikdache et al., 2020, Mikdache, Fontenas et al., 2020). *csp^-/-^* embryos injected with a constitutive active form of Rac1 were comparable to non-injected *csp^-/-^* embryos and showed a total lack of radial sorting and axonal ensheathment (Figure 8K,L).

This result suggests that *csp^-/-^* myelin defects do not ensue from a defective Rac1 activity.

### Forcing Laminin**α**2 expression fully rescues myelination defects in *csp^-/-^*

Previous studies have shown that Laminin211 promotes myelination via GPR126/cAMP dependent pathway (Petersen, Luo et al., 2015), and since forskolin treatment of *csp^-/-^* embryos is sufficient to re-establish myelination in these mutants, we hypothesized that Laminin 211 expression and/or function might be impaired in these mutants. To test this, we first analyzed Laminin expression at 48 hpf in controls and *csp^-/-^* embryos using immunostaining. Indeed, Laminin expression was significantly reduced along the PLLn of *csp^-/-^* embryos in comparison to controls (Figure 9A-C). We next wondered whether increasing extracellular Lama2 levels would rescue the myelination defect observed in *csp^-/-^*.

**Figure 9.**
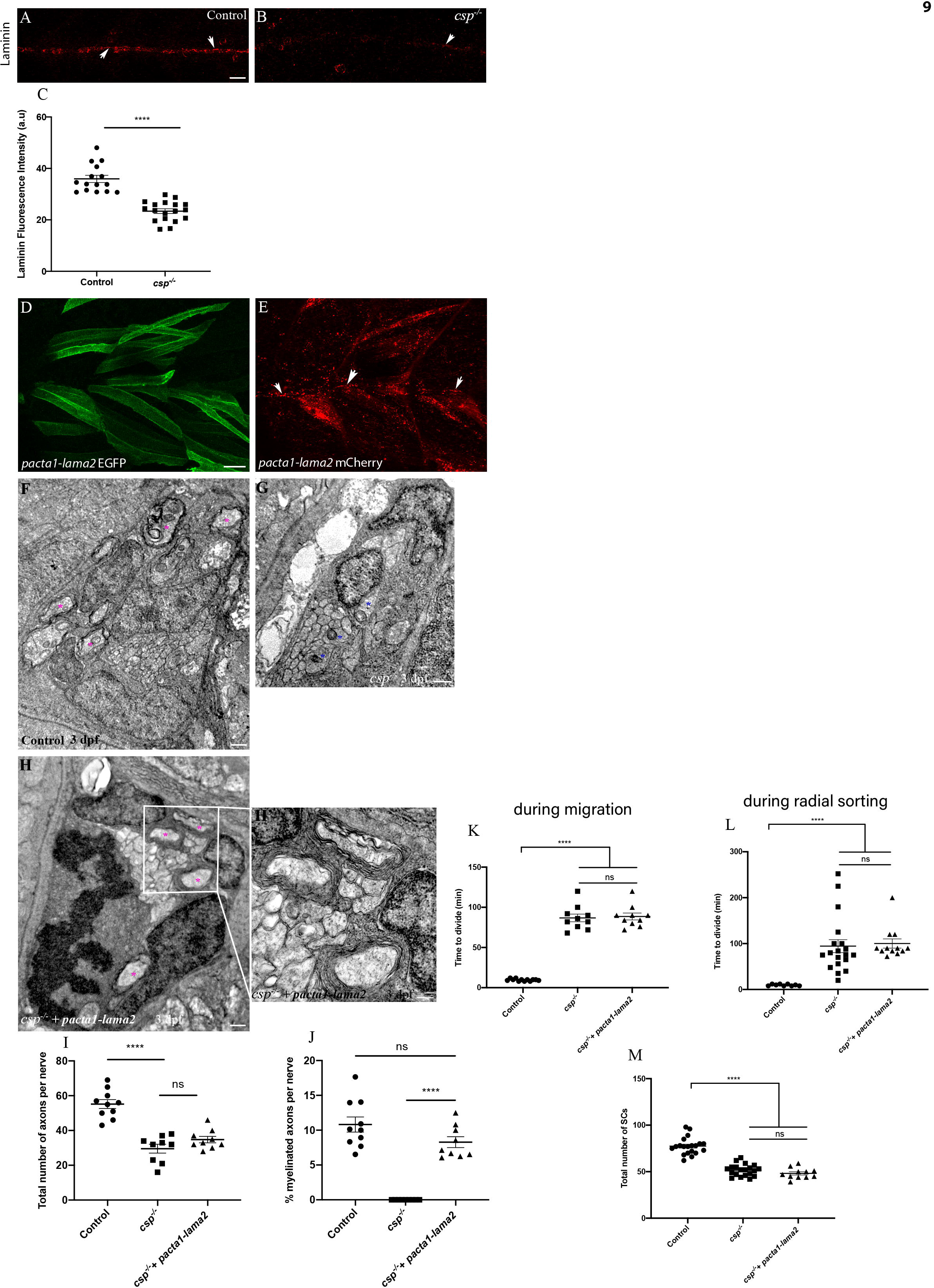
Laminin expression is significantly reduced in *csp^-/-^* and Laminin**α**2 over- expression rescues radial sorting and myelination defects in *csp^-/-^* embryos without increasing the number of Schwann cells Laminin expression in control (A, n=15) and *csp^-/-^* (B, n=18) embryos at 48 hpf showing the PLLn nerve (arrows). Scale bar = 20 μ . (C) Quantification of Laminin fluorescence intensity along the PLLn in controls (average of 35.95±1.39) and *csp^-/-^* (average of 23.37±0.90) at 48 hpf (****, p≤0.0001), a.u, arbitrary unit. (D) Lateral view of EGFP expression in muscles surrounding the PLLn following *pacta1-lama2* injection. Scale bar = 20 μm. (E) Lateral view of mCherry tagged secreted Laminin in muscles and within the PLLn (white arrows). TEM of a cross section of the PLLn at 3 dpf in control (F), *csp^-/-^* (G) and *csp^-/-^* + *pacta1-lama2* (H) embryos. Magenta asterisks represent some large caliber myelinated axons, some are shown at higher magnification in H’. Scale bars = 0.5 μm. (I) Quantification of total number of axons per nerve at 3 dpf in controls (average of 55.20±2.56, 10 nerves, n= 6 embryos), *csp^-/-^* (average of 29.56±2.55, 9 nerves, n= 7 embryos) and *csp^-/-^* + *pacta1-lama2* (average of 34.78±1.84, 9 nerves, n= 7 embryos) (****, p≤0.0001; ns, p=0.1190). (J) Quantification of the percentage of myelinated axons relative to the total number of axons per nerve at 3 dpf in controls (average of 10.82±1.085), *csp^-/-^* (average of 0±0) and *csp^-/-^* + *pacta1- lama2* (average of 8.28±0.77) (****, p≤0.0001; ns, p=0.075). (K) Quantification of the time required for control (average of 9.66±0.39, 12 cells, n= 4 embryos), *csp^-/-^* (average of 86.80±4.77, 10 cells, n= 4 embryos) and *csp^-/-^* injected with *pacta1-lama2*(average of 88.60±4.31, 10 cells, n= 4 embryos) Schwann cells to successfully complete mitotic division during migration (****, p ≤ 0.0001; ns, p=0.7828). min, minutes. (L) Quantification of the time required for control (average of 9.77±0.61, 9 cells, n= 4 embryos), *csp^-/-^* (average of 94.42±14.18, 19 cells, n= 4 embryos) and *csp^-/-^* injected with *pacta-lama2* (average of 100.2±10.01, 12 cells, n= 4 embryos) Schwann cells to successfully complete mitotic division during radial sorting (****, p ≤ 0.0001; ns, p=0.1424). (M) Quantification of the number of Schwann cells within a defined region of the PLLn at 48 hpf in controls (average of 77.45±2.05 cells, n= 20 embryos), *csp^-/-^* (average of 51.60±1.40, n= 20 embryos) and *csp^-/-^* injected with *pacta1-lama2* (average of 48.18±1.74 cells, n= 11 embryos), (****, p≤ 0.0001; ns, p=0.1402).

To this end, we took advantage of a *lama2* overexpression construct that expresses membrane EGFP as well as secreted mCherry-tagged Lama2 under the control of muscle-specific *skeletal-alpha actin (acta1)* promoter (Sztal, Sonntag et al., 2012) (Petersen et al., 2015) and injected 20 pg of this construct into *csp^-/-^* mutants. A wide and important amount of Laminin is secreted from muscle cells within the vicinity of PLLn in this case. Embryos that showed significant expression of EFGP within PLLn surrounding muscles and mCherry-tagged within PLLn (Figure 9D,E) were then selected and analyzed by TEM at 3 dpf. *csp^-/-^* injected embryos showed a significant increase in the number of myelinated axons in comparison to non-injected *csp^-/-^* larvae and were comparable to controls (WT *lama2* injected embryos) for the percentage of myelinated axons relative to the total number of axons (Figure 9F-J).

These results strongly suggest a defective Laminin expression and function within the PLLn of *csp^-/-^* mutant that is responsible for its radial sorting and myelination defect.

Laminin2 is known to control several aspects of Schwann cell development such as proliferation and differentiation (Chernousov et al., 2008). We therefore wondered whether Lama2 overexpression impacts on Schwann cell timely division and/or proliferation to mediate myelination or it just triggers the radial sorting/myelination process regardless of the number of Schwann cells. Indeed, Schwann cells numbers at 48 hpf were comparable between *csp^-/-^* and *csp^-/-^-pacta1-*lama2 injected embryos and showed a delay in mitotic exit similar to the one observed in *csp^-/-^* embryos during migration and prior to axonal ensheathment (Figure 9K-M). This result strongly suggests that the myelination defect observed in *csp^-/-^* is due to defective Laminin/cAMP pathway and that the temporal reduction in Schwann cells’ numbers is not essential for myelination to proceed.

### Schwann cells have a limited window of time to divide during migration in order to trigger myelination

Our findings have revealed an essential role for the temporal control of Schwann cell division during migration, mediated by Sil, in promoting radial sorting and myelination *in vivo* via Laminin2/cAMP pathway. This led us to hypothesize that Schwann cells might have a limited window of time during which they have to divide while migrating in order to trigger radial sorting and myelination once migration is completed. First, we tested whether blocking cell division during migration would alter Laminin expression within the PLLn as observed in *sil* mutants. Embryos incubated in aphidicolin during Schwann cell migration showed a significant decrease in the expression of Laminin along the PLLn at 48 hpf (Figure 10A-C). Second, we treated embryos with aphidicolin for different periods of time during migration and our results showed that Schwann cells have a limited time window of around 8 h in which they have to divide in order to myelinate (Figure 10F and S3; see Materials and Methods for aphidicolin treatment/efficacy). To test if this temporal delay in division that results in radial sorting and myelination defects is related to a defective Laminin expression/function, we injected embryos with 20 pg of *lama2* overexpression construct and treated embryos with aphidicolin between 22 and 30 hpf. The group of *lama2* injected embryos and treated with aphidicolin showed a significant increase in the number of myelinated axons in comparison to treated non-injected embryos (Figure 10D-H) and the percentage of myelinated axons relative to the total number of axons was similar to controls (Figure 10I).

**Figure 10.**
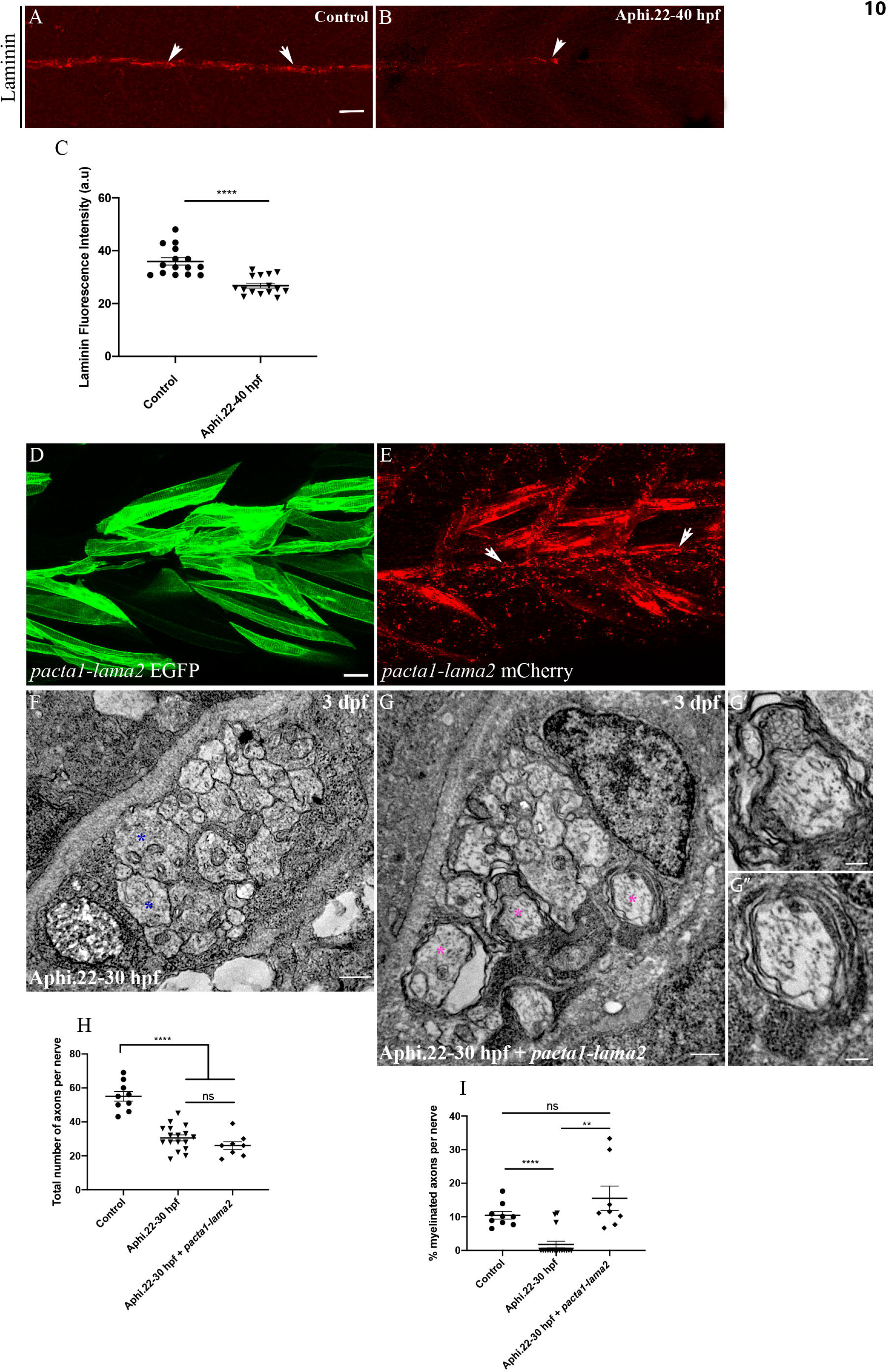
Laminin**α**2 overexpression rescues defective myelination in embryos with delayed Schwann cell division during migration Laminin expression in a control embryo (A, n=15) and aphidicolin treated embryo (B, n=15) at 48 hpf showing the PLLn nerve (arrows). Scale bar = 20 m. (C) Quantification of Laminin fluorescence intensity along the PLLn in controls (average of 36±1.4) and aphidicolin treated embryos (average of 26.80±0.92) at 48 hpf (****, p≤0.0001), a.u, arbitrary unit. (D) Lateral view of EGFP expression in muscles surrounding the PLLn following *pacta1-lama2* injection. Scale bar = 20 μm. (E) Lateral view of mCherry-tagged secreted Laminin in muscles and within the PLLn (white arrows). TEM of a cross section of the PLLn at 3 dpf in aphidicolin treated embryo (F) and aphidicolin treated embryo injected with *pacta1-lama2* (G). Magenta asterisks represent some large caliber myelinated axons, (some are shown at higher magnification in G’,G’’, scale bars = 0.2 μm). Blue asterisks represent some large caliber non myelinated axons. Scale bars = 0.5 μm. (H) Quantification of the total number of axons per nerve at 3 dpf in controls (average of 55.56±2.7), aphidicolin treated embryos (average of 30.53±1.75) and aphidicolin treated embryos + *pacta1-lama2* (average of 26.00±2.32) (****, p≤0.0001; ns, p=0.1401). (I) Quantification of the percentage of myelinated axons relative to the total number of axons per nerve at 3 dpf in controls (average of 10.73±1.4, 9 nerves, n= 6 embryos), aphidicolin treated embryos (average of 1.77±0.96, 17 nerves, n=13 embryos) and aphidicolin treated embryos + *pacta-lama2* (average of 15.53±3.61, 8 nerves, n=9 embryos) (****, p≤0.0001; **, p=0.0063; ns, p=0.2166).

This result strongly suggests a restricted time window for Schwann cells to divide while migrating during which Laminin expression is triggered.

### Schwann cells are a major source of Laminin within the PLL nerve and forcing Laminin**α**2 expression within Schwann cells rescues Mbp expression in *csp^-/-^* and aphidicolin treated embryos

Results so far have shown a strong correlation between timely Schwann cell division during migration, Laminin expression and subsequent myelination. However, it remained important to determine the source of Laminin within the PLLn and whether Schwann cell specific expression of Laminin would rescue myelination autonomously in *sil* mutant and aphidicolin treated embryos. To do so, we first incubated embryos with AG1478 that specifically blocks ErbB signaling in zebrafish and inhibits Schwann cell migration along the PLLn (Lyons et al., 2005). Embryos treated with AG1478 between 24 and 48 hpf presented no Schwann cells along the PLLn and showed a significant decrease in Laminin expression at 48 hpf (Figure 11A-C).

**Figure 11.**
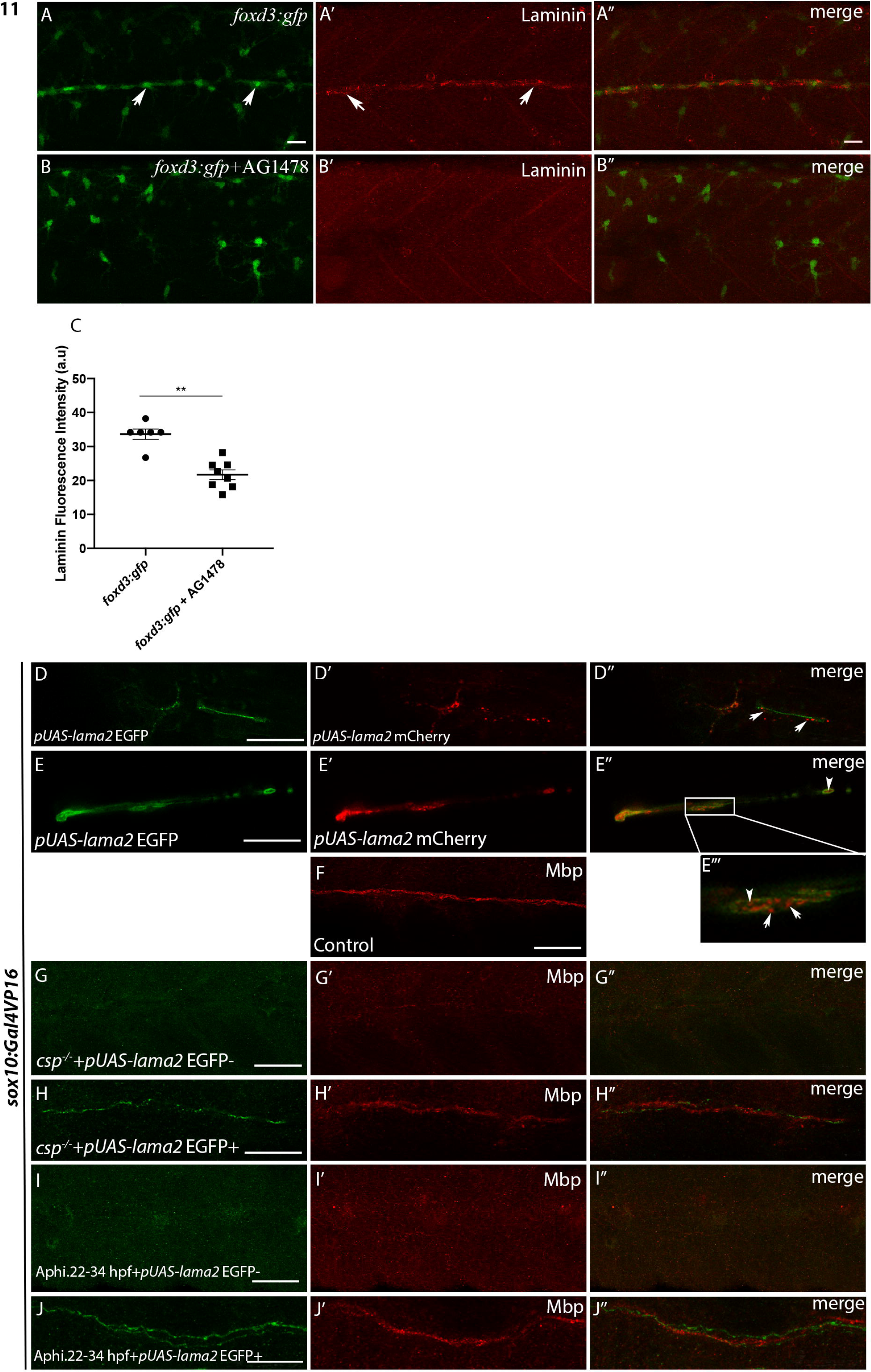
Laminin expression is significantly reduced in AG1478 treated embryos and Laminin**α**2 over-expression within Schwann cells rescues Mbp expression defects in *csp^- /-^* and aphidicolin treated embryos Laminin immuno-staining in *foxd3:gfp* (A-A’’, n=6) and *foxd3:gfp* embryos treated with AG1478 (B-B’’, n=8) at 48 hpf. Arrows indicate GFP+ Schwann cells in A and Laminin expression along the PLLn in A’. Scale bar = 20 μm. A’’ and B’’ are the merge images of A and A’ and of B and B’ respectively. (C) Quantification of Laminin fluorescence intensity along the PLLn in *foxd3:gfp* (average of 33.63±1.52) and *foxd3:gfp*/AG1478 treated embryos (average of 21.67±1.44) (**, p=0.0013), a.u, arbitrary unit. (D,E) Lateral views of EGFP expression in Schwann cells of the PLLn following *pUAS-lama2* injection in *sox10:Gal4VP16*. Scale bars = 20 μ (D’,E’) Lateral views of mCherry-tagged secreted Laminin in Schwann cells and within the PLLn. (D’’,E’’) merge of D and D’ and of E and E’ respectively. mCherry tagged secreted Laminin is highlighted in E’’’ at higher magnification; white arrows in D’’ and E’’’ indicate extracellular Laminin within the PLLn and white arrowheads in E’’ and E’’’ indicate mCherry tagged Laminin within Schwann cells. (F) Lateral view of Mbp immuno-labeling in control embryo at 3 dpf. Scale bar = 50 μm. (G, control, n=24) Lateral view showing the absence of EGFP expression in Schwann cells of the PLLn that correlates with a significant decrease in Mbp immuno-labeling (G’) following *pUAS-lama2* injection but with no EGFP/mCherry expression in *csp^-/-^/sox10:Gal4VP16*. Scale bar = 50 μm. (G’’) merge of G and G’. (H, n=8) Lateral view showing EGFP expression in Schwann cells of the PLLn that correlates with a normal Mbp immuno-labeling (H’) following *pUAS-lama2* injection with positive clones of EGFP/mCherry in *csp^-/-^ /sox10:Gal4VP16*. Scale bar = 20 μ . (H’’) merge of H and H’. (I, control, n=26) Lateral view showing the absence of EGFP expression in Schwann cells of the PLLn that correlates with a significant decrease in Mbp immuno-labeling (I’) following *pUAS-lama2* injection but with no EGFP/mCherry expression in *sox10:Gal4VP16* treated with aphidicolin between 22 and 34 hpf. Scale bar = 50 μm. (I’’) merge of I and I’. (J, n=9) Lateral view showing EGFP expression in Schwann cells of the PLLn that correlates with a normal Mbp immuno-labeling (J’) following *pUAS-lama2* injection with positive clones of EGFP/mCherry in *sox10:Gal4VP16* treated with aphidicolin between 22 and 34 hpf. Scale bar = 20 μm. (J’’) merge of J and J’.

This result establishes Schwann cells as a major source of Laminin within the PLLn.

Second, we designed a *pUAS-lama2-EGFP-mcherry* construct and took advantage of the *tg(sox10:Gal4VP16)* to allow a specific expression of Lama2 within Schwann cells (EGFP+ labeling); Lama2 is also found secreted within the PLLn (mCherry+ labeling) (Figure 11D- E’’’). We injected 20 pg of plasmid in *sox10:Gal4VP16* and *csp^-/-^/ sox10:Gal4VP16* at one cell stage and proceeded for EGFP and Mbp immunostaining for clonal analysis at 3 dpf. Injected *csp^-/-^* embryos that showed an expression of EGFP in Schwann cells along the PLLn expressed normal levels of Mbp while those that were negative to EGFP labeling showed a significant decrease in Mbp expression similar to non-injected *csp^-/-^* larvae (Figure 11F-H’’). Moreover, we injected *pUAS-lama2-EGFP-mcherry* construct in *sox10:Gal4VP16* and incubated embryos in aphidicolin during Schwann cell migration (22-34 hpf). Aphidicolin treated embryos that showed an expression of EGFP in Schwann cells along the PLLn expressed normal levels of Mbp while those that were negative to EGFP labeling showed a significant decrease in Mbp expression at 3 dpf (Figure 11I-J’’).

These results strongly suggest that Schwann cell division during migration is largely responsible of early Laminin expression within the PLLn. This constitutes a crucial step for timely myelination.

## Discussion

Schwann cells migrate and divide along peripheral axons and the intimate contact between the two contributes to Schwann cell proliferation, mainly through the neuregulin/ErbB axis. Other signals provided by ECM proteins that Schwann cells secrete, are also important for their proliferation (Bunge, Clark et al., 1990, Bunge, Williams et al., 1982, Lyons et al., 2005, Michailov, Sereda et al., 2004, Nave & Salzer, 2006). Here, we investigate the characteristics of Schwann cell division during migration and radial sorting and their impact on myelination. We show that the temporal control of mitotic exit within Schwann cells is tightly controlled by the spindle pole protein Sil. We provide evidence that timely division of Schwann cells during migration is essential for radial sorting and myelination.

Schwann cells must regulate their numbers in order to myelinate or ensheath all axons that are supposed to be myelinated by the end of radial sorting (Jessen & Mirsky, 2005), so one would predict that reducing the number of Schwann cells in the nerve would result in a reduction of myelinated axons. Indeed, several molecules that control Schwann cell proliferation have been identified such as Cdc42, focal adhesion kinase and several extracellular matrix proteins of the laminin family such as laminin-2, 8 and γ1 (Benninger, Thurnherr et al., 2007, Grove, Komiyama et al., 2007, Yang, Bierman et al., 2005). Specific ablation of these molecules in Schwann cells causes a reduction in the number of myelinated axons.

However, how important is the timely division of Schwann cells? If Schwann cells divide intensively during radial sorting to generate the right numbers prior to axonal ensheathment, then how would early division during migration contribute to myelination? Are insufficient Schwann cell numbers *per se* a cause for radial sorting and myelination defects? The importance of Schwann cell division during radial sorting has been examined *in vivo* (Raphael et al., 2011), showing that blockade of cell division leads to significant defects in radial sorting and myelination by Schwann cells. Several parameters should be considered in order to assess the role of division in myelination following aphidicolin treatment, that is obviously the drastic decrease in the total number of axons in the PLLn and the reduction in the number of Schwann cells. Our results show that Schwann cell division during radial sorting is not required for myelination *per se* since blocking cell division reduces the number of myelinated axons in the PLLn but not the ratio of myelinated axons relative to the total number of axons resulting from this treatment. While aphidicolin-treated embryos during radial sorting show a normal percentage of myelinated axons, *csp^-/-^* ones are totally devoid of radial sorting and myelination. However, when analyzing earlier events, we discovered that early Schwann cell division during migration, as well as the temporal control of its mitotic exit are essential in this process. These findings demonstrate that Schwann cells have a restricted time window during which they have to divide, specifically while migrating, in order to set up and initiate myelin wrapping at later stages.

How might Sil control radial sorting and myelination in the PNS?

Since *csp^-/-^* embryos show a radial sorting defect, it remains possible that the myelination defects result, at least partially, from a defective cytoskeletal re-arrangement mediated by Rac1. Indeed, Schwann cells require a drastic process extension of their membrane in order to sort individual axons from within a bundle. A group of genes that regulate this aspect of Schwann cell behavior has been identified and includes *elmo1*, *dock1, β1 integrin* and *ILK* (Cunningham, Herbert et al., 2018, Feltri, Graus Porta et al., 2002, Mikdache et al., 2020).

Some evidence points towards a role for these molecules in activating the small GTPase Rac1 that acts upon Schwann cells cytoskeleton during radial sorting (Guo, Moon et al., 2012, Nodari et al., 2007). Our analysis strongly suggests that the myelination defects observed in *csp*^-/-^ do not result from a defective Rac1 activity since forcing the expression of Rac1 does not rescue, even partially, this defect.

Schwann cells in *csp*^-/-^ struggle to progress through the M phase of the cell cycle and show a severe reduction in the expression of *krox20*. This phase is characterized by chromatin condensation and by ejection of transcription factors and chromatin binding proteins (Ma et al., 2015). Schwann cells are no exception since major chromatin remodeling complexes are present and play an important role in temporal Schwann cell development (Frob & Wegner, 2019, Gomis-Coloma, Velasco-Aviles et al., 2018). It is possible that Schwann cell transcription factors required for the initiation of myelination are permanently ejected in *csp*^-/-^ following the temporal defect in M phase progression and therefore unable to bind the DNA.

Another possibility is a defective upstream signaling activity in *csp^-/-^* embryos that blocks Schwann cell differentiation, assuming that chromatin accessibility is re-established after division. Our analysis suggests that it is rather a defective signaling activity that is responsible for the peripheral lack of myelin observed in *csp*^-/-^ embryos and that transcription activity is responsive to upstream signaling cues.

Two major signaling pathways are required for several aspects of Schwann cell development, that are the Neuregulin/ErbB and the Gpr126/cAMP signaling pathways (Monk et al., 2009, Newbern & Birchmeier, 2010). The neuregulin/ErbB plays a major role in migration, survival, proliferation, radial sorting and myelination by Schwann cells through a plethora of downstream factors such as PI3K/Akt, MEK/ERK and Cdc42 (Glenn & Talbot, 2013, Newbern & Birchmeier, 2010). The Gpr126 pathway signals through cAMP and activates myelination by elevating the levels of cAMP so that a forskolin treatment of *gpr126* mutants fully restores myelin ensheathment (Monk et al., 2009). However, elevating cAMP in zebrafish *erbb2* mutants does not rescue its myelin defects (Glenn & Talbot, 2013). A forskolin treatment of *csp*^-/-^ embryos fully rescues radial sorting defects, restores *krox20* and *mbp* expression as well as axonal ensheathment, suggesting a defective cAMP pathway activity in these mutants.

A particularly interesting observation is the difference in *sil* deficiency phenotype between retina and Schwann cells. Sil is a ubiquitously expressed protein and is involved in the fundamental process of mitosis but is highly enriched in the nervous system. Neurons might in this case be more sensitive to Sil defects leading to massive apoptosis within the central nervous system (CNS); however, it is Schwann cell differentiation that is impaired in *csp*^-/-^ with no increase in apoptosis. It is therefore possible that different cells might respond differently to *sil* defects and our results suggest that MCPH genes have a broader requirement during development that is not solely confined to neuronal cells in the CNS. It is also important to note that zebrafish dynein cytoplasmic 1 heavy chain 1 (*dync1h1*) mutant shows comparable myelin defects to *csp*^-/-^ (Langworthy & Appel, 2012). Dynein is involved in spindle organization and checkpoint (Grava, Schaerer et al., 2006, Kiyomitsu & Cheeseman, 2012, Silva, Reis et al., 2014) similar to Sil and therefore it is highly possible that spindle checkpoints proteins are fundamentally required for the temporal control of mitosis within Schwann cells in order to initiate myelination.

One important question remained to be solved: what is the molecular link between Schwann cell division, its temporal control and cAMP pathway? Our analyses have shown an essential role for Sil in Schwann cell differentiation, this added to the fact that cAMP stimulation rescued the myelination defects prompted us to test for upstream signals that mediates cAMP activity. Several studies have now established an essential role for the Gpr126 in driving cAMP activity in Schwann cells through G proteins (Mogha, Benesh et al., 2013) and this has recently been attributed to Gpr126 interaction with ECM, more specifically with Laminin211 (Petersen et al., 2015). It is therefore possible that *sil* deficient Schwann cells are incapable of secreting extracellular proteins and/or responding to them to drive the ECM/Gpr126/cAMP pathway (Paavola, Sidik et al., 2014, Petersen et al., 2015). Our results showed that *csp^-/-^* larvae presented a significant decrease in Laminin expression in the PLLn and forcing Laminin2 expression within the vicinity of the nerve was sufficient to activate the myelination program in Schwann cells, drive radial sorting and re-establish axonal wrapping.

Based on all these findings, it was therefore interesting to assess the temporal setting and the mechanism by which Schwann cell division triggers myelination. This study and others have shown that Schwann cells go through extensive cell proliferation prior to myelination at different timepoints of their development: (i) during migration and (ii) during radial sorting once they finished migrating in zebrafish (Lyons et al., 2005) and rodents (Stewart, Morgan et al., 1993). To understand how timely division impact on Schwann cell differentiation, we used the cell division inhibitor, aphidicolin, to temporally block cell division at these two essential timepoints of intensive Schwann cell proliferation and analyzed its impact on myelination. To our surprise, it was Schwann cell division during migration that dictated the radial sorting and myelination events. In addition to characterizing how temporal control of mitotic exit impacts myelination, our study also provided documentation of the existence of a restricted window of time (around 8 hours) during which Schwann cells have to divide while migrating in order to initiate radial sorting and myelination *in vivo*. Several studies have pointed to a role for Laminin in controlling Schwann cell proliferation but this is the first study that implicates Schwann cell division, particularly during migration, in Laminin expression and subsequent myelination. The temporal control of this particular division is therefore an essential step towards initiating the myelination program through Laminin/cAMP pathway, a role that stretches beyond simply increasing the pool of Schwann cells required to continuously engulf the nerve as it grows.

To come back to the question of numbers, is it important to generate enough Schwann cells to myelinate? Our findings reveal that the myelination defects observed in *csp^-/-^* and aphidicolin- treated embryos do not ensue from reducing Schwann cells numbers *per se*, since Laminin overexpression can fully rescue the percentage of myelinated axons within the PLLn even if Schwann cell numbers remain significantly reduced. It would be interesting to know in the future to what extent we can reduce the number of Schwann cells before they fully stretch along the axons and whether they would still be able to myelinate in presence of Laminin under acute mechanical strain.

Finally, we show that Schwann cells are a major source of Laminin within peripheral nerves in zebrafish. Our clonal analysis shows that forcing Laminin expression autonomously within Schwann cells is enough to secrete Laminin in the extracellular space and restore normal Mbp expression within the vicinity of these cells in *csp^-/-^* and aphidicolin treated embryos.

In summary, our work provides evidence of a novel ON/OFF mechanism that links early temporal control of Schwann cell division during migration to radial sorting and axonal wrapping via Laminin/cAMP pathway *in vivo*.

## Materials and Methods

### Embryo care

Embryos were staged and cared for according to standard protocols (https://zfin.org/zf_info/zfbook/cont.html). *Tg(foxd3:gfp)* stable transgenic line, that labels SCs was used in this study (Gilmour et al., 2002). *Tg(sox10:Gal4VP16)^el159^* was kindly provided by Gage Crump. *Csp^+/-^* embryos (*stil^cz65^)* were purchased from ZIRC. All animal experiments were conducted with approved protocols at Inserm by DDPP Val de Marne, France under license number F 94-043-013.

### Plasmid constructs

*pTol2-Sox10:Sil-P2A-mCherry:* The sil-P2A-mCherry cassette allowing simultaneous expression of sil and membrane-localized mCherry separated by the self-cleaving P2A peptide was generated by PCR amplification.

The 5′-CGATTCACTAGTATGAACCGTGTACAAGTGGATTTTAAAGG-3′ forward and 5′-GAAGTTCGTGGCTCCGGATCCAAAGAGCTTGGGGAGCTGCCGTAACC - 3′ reverse primers were used onto the pDNR-lib-STIL cDNA bought from addgene and the 5′- GGTTACGGCAGCTCCCCAAGCTCTTTGGATCCGGAGCCACGAACTTC-3′ forward and 5′-CTATGACCATGATTACGCCAAG-3′ reverse primers onto the pCS2-mCherry- CAAX plasmid.

The resulting PCR fragment was digested by SpeI and NotI before sub-cloning into the - 4725Sox10-cre vector (a gift from Robert Kelsh) to obtain the *pSox10:sil-P2A-mCherry* plasmid.

*p5UAS: lama2-mcherry-T2A-EGFPcaax :* The pT2i5uasbiH2bChe-GFP (a gift from Michel Volovitch) was used to amplify the 5 UAS sequence using 5’GTGGGCCCTGCGTCTAGAGTC3’ forward and 5’ CTCACCATATGGGCGACCGGTGG3’ reverse primers. Then, in order to replace *Acta1* promoter, the PCR product was subcloned into the *pacta1:lama2-mcherry-T2A- EGFPcaax* vector (a gift from Peter Currie and Kelly Monk) digested by ApaI and NdeI enzymes.

### Microinjections

*Mito:gfp* (a gift from Dominik Paquet) and *h2b ::gfp* (a gift from Jon Clarke) mRNAs were synthesized using SP6 mMessage mMachine System after linearization with Not1 and injected at 200 pg per embryo. *pTol2:sox10-sil-2A-mcherry-Caax* was injected at 10 pg per embryo along with 50 pg of Tol2 transposase mRNA (a gift from David Lyons). *Rac1V12* (constitutively active Rac1, kindly provided by Nicolas David) mRNA were synthesized using SP6 mMessage mMachine System after linearization with Not1 and injected at 2-10 pg per embryo. *pacta1-lama2* (kindly provided by Kelly Monk and Peter Currie) was injected at 20 pg per embryo along with 100 pg of Tol2 transposase mRNA. *pUAS-lama2* was injected at 20 pg per embryo.

### In situ hybridization

*In situ* hybridization was performed following standard protocols using *krox20* and *mbp* probes (Fontenas, De Santis et al., 2016). Embryos were raised in egg water with PTU (0.003%) to avoid pigmentation and were fixed overnight in 4% paraformaldehyde at 52 hpf and 3 dpf. Embryos were dehydrated the next day and kept in methanol at -20°C. Embryos were then rehydrated and treated with proteinase K and incubated overnight with the corresponding probe. Embryos were then washed and the staining was revealed using anti-digoxigenin antibody.

### Immunofluorescence

The following antibodies and dilutions were used: mouse anti-acetylated tubulin (Sigma; 1/500; T7451), mouse anti-HuC/HuD neuronal protein (Invitrogen; 1/500; REF A21271), rabbit anti-laminin (Sigma, 1/200; L9393), mouse anti-phospho Histone 3 (Ser 10) (Millipore; 1/500; Cat # 06-570), rabbit anti-myelin basic protein (Custom produced (Tingaud-Sequeira Angèle , Anja Knoll-Gellida a et al., 2017)), gift from Patrick Babin, 1/500), mouse anti-EGFP (Millipore; 1/500; REF MAB3580). Primary antibodies were detected with appropriate secondary antibodies conjugated to either Alexa 488, Alexa 568 or Alexa 647 (Invitrogen) at a 1:500 dilution. For immunostaining, embryos were fixed in 4% paraformaldehyde 1X PBS overnight at 4°C, then washed with 1xPBS to be dehydrated in Methanol for at least 6 hours at -20°C. Samples were then rehydrated, digested with proteinase K and blocked with 0.5% triton in PBS and 10% sheep serum and then incubated with primary antibody overnight at 4°C (diluted in PBS+2% sheep serum). Larvae were then washed with PBS for a few hours and then incubated with secondary antibody in 0.5% triton in PBS and 2% sheep serum overnight at 4°C. Stained larvae were then imaged with a Leica SP8 confocal microscope. For Laminin and Mbp staining, embryos were fixed in 4% PFA for 30 min and treated for 7 min with acetone at -20°C. Samples were then blocked with 0.5% triton in PBS and 10% sheep serum and then incubated with primary antibody overnight at 4°C (diluted in PBS+2% sheep serum). Larvae were then washed with PBS for a few hours and then incubated with secondary antibody in 0.5% triton in PBS and 2% sheep serum for 3 hours at RT. For Laminin fluorescence intensity quantification, the same parameters (Excitation/emission, gain for detectors, lasers intensity) were applied for images’ acquisition in controls, *csp^-/-^* and aphidicolin treated embryos and fluorescence intensity was measured using image J (Analyze/measure).

### Aphidicolin, forskolin and AG1478 treatment

Embryos were incubated in fish water containing 150 μM aphidicolin diluted in DMSO between 45 and 54 hpf or 45 and 72 hpf and with 100 μM between 22 and 40 hpf, 22 and 30 hpf or 22 and 34 hpf. Controls were incubated in equivalent amount of DMSO (0.5 and 0.33%) solution during the same periods. It took around 2 to 3 h for the aphidicolin to become fully efficient following incubation.

Embryos were incubated in fish water containing 20 μM forskolin diluted in DMSO between 45 and 54 hpf. Controls were incubated in equivalent amount of DMSO solution (0.1%) during the same period.

Embryos were incubated in fish water containing 4 μM AG1478 diluted in DMSO between 24 and 48 hpf. Controls were incubated in equivalent amount of DMSO solution (0.1%) during the same period.

### Acridine orange staining

Embryos were anesthetized with 0.03% tricaine and incubated in fish water containing 5 μM acridine orange for 20 minutes in the dark. After wash, they were embedded in 1.5% low melting point agarose, and imaged with a Leica SP8 confocal microscope.

### Live imaging

Embryos were anesthetized with 0.03% tricaine and embedded in 1.5% low melting point agarose. For mito:GFP tracking experiments, PLLn was examined at 48 hpf from a lateral view. A series of 10 minutes time-lapses were recorded. Recordings were performed at 27°C using a Leica SP8 confocal microscope. For *Tg(foxd3:gfp), Tg(foxd3:gfp)/csp^-/-^* as well as controls and *csp^-/-^* injected with *h2b:gfp*, PLLg was examined at 28 hpf and 48 hpf and recordings were acquired for up to 12h.

### Transmission electron microscopy

At 3 and 4 dpf, embryos were fixed in a solution of 2% glutaraldehyde, 2% paraformaldehyde and 0.1M sodium cacodylate pH 7.3 overnight at 4°C. This was followed by a post-fixation step in cacodylate-buffered 1% osmium tetroxide (OsO_4_, Serva) for 1h at 4°C and in 2% uranyl acetate for 1h at room temperature. The tissue was then dehydrated and embedded in epoxy resin. Sections were contrasted with saturated uranyl acetate solution and were examined with a 1010 electron microscope (JEOL) and a digital camera (Gatan).

### Statistical analysis

Means and standard deviations were calculated with Graph Pad Prism 7. All data were first tested for normal distribution using the ShapiroWilk normality test combined with D’Agostino & Pearson normality test. All experiments with only two groups and one dependent variable were compared using an unpaired *t*-test with Welch’s correction if they passed normality test; if not, groups were compared with the nonparametric Mann-Whitney test. Statistically significant differences were determined using one-way ANOVA for all experiments with more than two groups but only one dependent variable. Error bars depict standard errors of the mean (SEM). ns, p>0.05; *, p≤0.05; **, p≤0.01; ***, p≤0.001; ****, p 0.0001. n represents the number of embryos.

## Acknowledgments

We would like to thank Peter Currie and Kelly Monk for the *pacta1-lama2* plasmid, Michel Volovitch and Sophie Vriz for *pUAS* construct and discussing cloning strategy, Robert Kelsh for *sox10* promoter, Gage Crump for *Tg(sox10:Gal4VP16)^el159^* embryos, Patrick Babin for providing Mbp antibody and Jon Clarke for his critical reading of the manuscript. This work was funded by Inserm and University Paris-Saclay. Emilie Lesport is funded by “Institut Professeur Baulieu”.

## Supplemental information

Figure S1. Timeline of Schwann cell behavior and myelination in zebrafish. Schwann cells migrate and divide along growing axons between 24 hpf and 48 hpf. They start to radially sort axons of the PLLn at around 48 hpf and divide intensively. Myelin sheath is analyzed starting from 3 dpf (or 72 hpf). hpf, hours post fertilization; dpf, days post fertilization; PLLn, posterior lateral line nerve.

Figure S2. Sil is not required for axonal growth nor for mitochondrial axonal transport along the PLLn but regulates the number of neurons within the PLLg

(A) Acetylated tubulin expression in a control embryo (n=12) and *csp^-/-^* embryo (n=13) at 48 hpf showing the PLLn nerve (arrows). Lateral views of a control (n=16) and *csp^-/-^* embryo (n=11) at 48 hpf showing PLLn GFP-expressing Schwann cells (arrows). Scale bars = 50 μ

(B) Still images from time-lapse imaging in control and *csp^-/-^* embryos injected with *mito:gfp*. Arrows point to the same mitochondria followed through time in control and *csp^-/-^*. Scale bar = 5 μm. s, seconds.

(C) Quantification of the average speed of mitochondria along the PLLn at 50 hpf in controls (302 mitochondria, n= 8 embryos) and *csp^-/-^* embryos (93 mitochondria, n= 6 embryos) (ns, p=0.6976).

(D) HuC immuno-labeling of the PLLg at 48 and 72 hpf in control and *csp^-/-^* embryos. Scale bar = 5 μm.

(E) Quantification of the number of neurons in the PLLg at 48 hpf in control (average of 54.58±5.99 neurons, n= 12) and *csp^-/-^* (average of 31.71±5.31 neurons, n= 7) embryos and at 72 hpf in control (average of 74.33±5.88 neurons, n= 12) and *csp^-/-^* (40.29±1.60 neurons n= 7) embryos (****, p≤0.0001; ****, p≤0.0001).

Figure S3. Schwann cells have a limited window of time to divide during migration in order to radially sort axons and myelinate

TEM of a cross section of the PLLn at 5 dpf in control (A) and embryo treated with aphidicolin between 22 and 34 hpf (B). Magenta asterisks highlight some large caliber myelinated axons (some shown at higher magnification in A’, scale bar = 0.2 μm) and blue asterisks show some large caliber non-myelinated axons (some shown at higher magnification in B’, scale bar = 0.2 μm). Scale bars = 0.5 μm. (C) Quantification of the number of myelinated axons per nerve at 5 dpf in controls (average of 9±1.29, 4 nerves, n= 4 embryos) and embryos treated with aphidicolin between 22 and 34 hpf (average of 0, 4 nerves, n= 4 embryos) (**, p=0.0061).

(D) Quantification of the total number of axons per nerve at 5 dpf in controls (average of 49.75±3.32) and embryos treated with aphidicolin between 22 and 34 hpf (average of 28.75±3.27) (**, p=0.0041).

(E) Quantification of the percentage of myelinated axons relative to the total number of axons per nerve at 5 dpf in controls (average of 17.94±1.92) and embryos treated with aphidicolin between 22 and 34 hpf (average of 0) (**, p=0.0021).

(F) Quantification of the percentage of axons according to their diameter relative to the total number of axons per nerve at 5 dpf in controls (average of 66.66±5.26 for 0-0.4 μm; average of 33.34±5.27 for >0.4 μm) and aphidicolin treated embryos between 22 and 34 hpf (average of 54.34±6.7 for 0-0.4 μm; average of 45.66±6.75 for >0.4 μ). ns, p=0.2209.

Movie S1. Real-time imaging of Schwann cells in *Tg(foxd3:gfp)* at 48 hpf.

A 48 hpf embryo expressing GFP in Schwann cells; the control embryo was imaged every 4 minutes for several hours by confocal microscopy. Lateral view; anterior to the left and dorsal to the top. This video represents four and a half hours of continuous real-time imaging.

Movie S2. Real-time imaging of Schwann cells in *Tg(foxd3:gfp)/ csp^-/-^* at 48 hpf.

A 48 hpf embryo expressing GFP in Schwann cells; the *csp^-/-^* embryo was imaged every 4 minutes for several hours by confocal microscopy. Lateral view; anterior to the left and dorsal to the top. This video represents four and a half hours of continuous real-time imaging.

Movie S3. Real-time imaging of mitochondria in a control PLLn at 48 hpf.

A 48 hpf control embryo expressing GFP in mitochondria after *mito-gfp* mRNA injection; the embryo was imaged every 120 milliseconds for several minutes by confocal microscopy. Lateral view; anterior to the left and dorsal to the top. This video represents 18 seconds of real-time continuous imaging.

Movie S4. Real-time imaging of mitochondria in a *csp^-/-^* PLLn at 48 hpf.

A 48 hpf *csp^-/-^* embryo expressing GFP in mitochondria after *mito-gfp* mRNA injection; the embryo was imaged every 120 milliseconds for several minutes by confocal microscopy. Lateral view; anterior to the left and dorsal to the top. This video represents 36 seconds of continuous real-time imaging.

Movie S5. Real-time imaging of Schwann cell nuclei in control at 48 hpf.

A 48 hpf control embryo expressing GFP in Schwann cell nuclei after *h2b-gfp* mRNA injection; the embryo was imaged every 4 minutes for several hours by confocal microscopy. Lateral view; anterior to the left and dorsal to the top. This video represents 40 minutes of continuous real-time imaging.

Movie S6. Real-time imaging of Schwann cell nuclei in *csp^-/-^* at 48 hpf.

A 48 hpf *csp^-/-^* embryo expressing GFP in Schwann cell nuclei after *h2b-gfp* mRNA injection; the embryo was imaged every 4 minutes for several hours by confocal microscopy. Lateral view; anterior to the left and dorsal to the top. This video represents 100 minutes of continuous real-time imaging.

Movie S7. Real-time imaging of Schwann cells in *Tg(foxd3:gfp)* at 28 hpf

A 28 hpf embryo expressing GFP in Schwann cells; the control embryo was imaged every 4 minutes for several hours by confocal microscopy. Lateral view; anterior to the left and dorsal to the top. This video represents two hours of continuous real-time imaging.

Movie S8. Real-time imaging of Schwann cells in *Tg(foxd3:gfp)/ csp^-/-^* at 28 hpf.

A 28 hpf embryo expressing GFP in Schwann cells; the *csp^-/-^* embryo was imaged every 4 minutes for several hours by confocal microscopy. Lateral view; anterior to the left and dorsal to the top. This video represents four hours of continuous real-time imaging.

Movie S9. Real-time imaging of Schwann cells in *Tg(foxd3:gfp)* at 48 hpf.

A 48 hpf embryo expressing GFP in Schwann cells; the control embryo was imaged every 3 minutes for several hours by confocal microscopy. Lateral view; anterior to the left and dorsal to the top. This video represents three hours of continuous real-time imaging.

Movie S10. Real-time imaging of Schwann cells in *Tg(foxd3:gfp)* treated with aphidicolin between 22 and 40 hpf at 48 hpf.

A 48 hpf embryo expressing GFP in Schwann cells; the aphidicolin treated embryo was imaged every 3.5 minutes for several hours by confocal microscopy. Lateral view; anterior to the left and dorsal to the top. This video represents six hours of continuous real-time imaging.

